# scINSIGHT for interpreting single-cell gene expression from biologically heterogeneous data

**DOI:** 10.1101/2021.10.13.464306

**Authors:** Kun Qian, Shiwei Fu, Hongwei Li, Wei Vivian Li

## Abstract

The increasing number of scRNA-seq data emphasizes the need for integrative analysis to interpret similarities and differences between single-cell samples. Even though different batch effect removal methods have been developed, none of the existing methods is suitable for heterogeneous single-cell samples coming from multiple biological conditions. To address this challenge, we propose a method named scINSIGHT to learn coordinated gene expression patterns that are common among or specific to different biological conditions, offering a unique chance to identify cellular identities and key biological processes across single-cell samples. We have evaluated scINSIGHT in comparison with state-of-the-art methods using simulated and real data, which consistently demonstrate its improved performance. In addition, our results show the applicability of scINSIGHT in diverse biomedical and clinical problems.

## Introduction

Single-cell RNA sequencing (scRNA-seq) technologies enable gene expression measurement at a single-cell resolution, and have opened a new frontier to understand animal development, physiology, and disease-associated molecular mechanisms [1, 2, 3, 4, 5]. Rapid advances of scRNA-seq technologies have resulted in the generation of large-scale single-cell gene expression datasets from different platforms in different laboratories [6, 7], using samples that span a broad range of species, tissue types, and experimental conditions [8, 9, 10]. The increasing number of scRNA-seq datasets emphasizes the need for integrative biological analysis to help assess and interpret similarities and differences between single-cell samples and to obtain in-depth insights into the underlying biological systems [11, 12, 13]. For example, integrative analysis of human and mouse transcriptomes has identified conserved cell types and transcription factors in pancreatic cells [14]; integrative analysis of scRNA-seq data from multiple melanoma tumors has identified a resistance program in malignant cells that is associated with T cell exclusion and immune evasion [15].

A fundamental goal in integrative scRNA-seq data analysis is to jointly define cell clusters, obtain their functional interpretation and annotation, and identify differentially activated biological pathways in distinct cell types and biological conditions. However, a key challenge for achieving this goal is the heterogeneity present in single-cell gene expression data. As expression data from different sources are associated with various types of technical effects [16], expression patterns of biological interest need to be discerned from cell-specific and sample-specific effects in order to compare single-cell transcriptomes across samples and biological contexts. In addition to technical variability, genuine cellular heterogeneity is present in different cell types and cell states with distinct behaviors and functions, and in response to different perturbations [17].

To help remove the batch effects emerging from scRNA-seq data generated by different sequencing platforms or library-preparation protocols, several batch correction methods, including mnnCorrect [18], BBKNN [19], and BEER [20], have been developed. However, batch correction methods assume that the differences between the single-cell samples are purely technical and non-biological, and thus are not appropriate for analyzing biologically different scRNA-seq datasets, such as tissue biopsy data from different patients [21, 22] or data of the same tissue type from related species [14, 23]. In practice, there are multiple integration methods that have been used to analyze single-cell gene expression data from biologically heterogeneous sources [24, 25, 26, 27, 28, 29]. For example, Seurat [24] matches cell states across samples by identifying the so-called anchor cells in a lower-dimensional space constructed with canonical correlation analysis. Similarly, Scanorama [25] matches cell clusters by identifying mutual nearest neighbors in a lower-dimensional space constructed with randomized singular value decomposition. scMerge [26] performs clustering in each sample, matches clusters across samples, and then uses control genes to correct for inter-sample variation. In addition, LIGER [27] identifies both shared and dataset-specific metagene factors to enable integration of multiple single-cell samples.

Even though the above methods have been shown useful in batch-effect removal and integrative analysis of multiple single-cell samples [30], they do not account for the situation where heterogeneous samples come from distinct biological conditions (e.g., different experimental groups or disease phases), and thus may compromise the results. To address this challenge, we propose a novel method named scINSIGHT (<u>IN</u>terpreting single cell gene expres<u>SI</u>on from biolo<u>G</u>ically <u>H</u>eterogeneous da<u>T</u>a) to jointly model and interpret gene expression patterns in single-cell samples from biologically heterogeneous sources. scINSIGHT uses a new model based on nonnegative matrix factorization (NMF) [31] to decompose gene expression patterns of distinct cell types and biological conditions. Compared with existing tools, scINSIGHT has the following advantages: (1) it explicitly models coordinated gene expression patterns that are common among or unique to biological conditions, enabling the decomposition of common and condition-specific gene modules from high-dimensional gene expression data; (2) it achieves precise identification of cell populations across single-cell samples, using common gene modules that capture cellular identities; (3) it enables efficient comparison between samples and biological conditions based on cellular compositions and module expression; (4) it discovers sparse and directly interpretable module expression patterns to assist functional annotation. We evaluated the performance of scINSIGHT in both simulation and real data studies, both of which demonstrated its accuracy and effectiveness for interpreting single cell gene expression from biologically heterogeneous data.

## Results

### scINSIGHT jointly models heterogeneous scRNA-seq datasets through matrix factorization

We propose a novel matrix factorization model named scINSIGHT to jointly analyze multiple single-cell gene expression samples that can be grouped based on biological conditions, such as different disease phases, treatment groups, or developmental stages. To the best of our knowledge, scINSIGHT is the first integration method for scRNA-seq data which can directly account for group information. It assumes that each gene module is a sparse and non-negative linear combination of genes, and each cell is jointly defined by the expression of common and condition-specific modules. Given multiple gene expression samples from different biological conditions, scINSIGHT aims to simultaneously identify common and condition-specific gene modules and quantify their expression levels in each sample in a lower-dimensional space (Figure 1A, Methods). To achieve joint matrix factorization, we construct an objective function which aims to minimize the factorization error, with constraints on the scale of the condition-specific components and the similarity between condition-specific gene modules. We propose an algorithm based on block coordinate descent [32] to obtain the solutions of the above optimization problem, and we also provide approaches to select parameters in the scINSIGHT model (Methods). With the factorized results, the inferred expression levels and memberships of common gene modules can be used to cluster cells and detect cellular identities (Figure 1B), the condition-specific gene modules can help compare functional differences in transcriptomes from distinct conditions (Figure 1C), and the reconstruction residuals are treated as technical noises.

**Figure 1:**
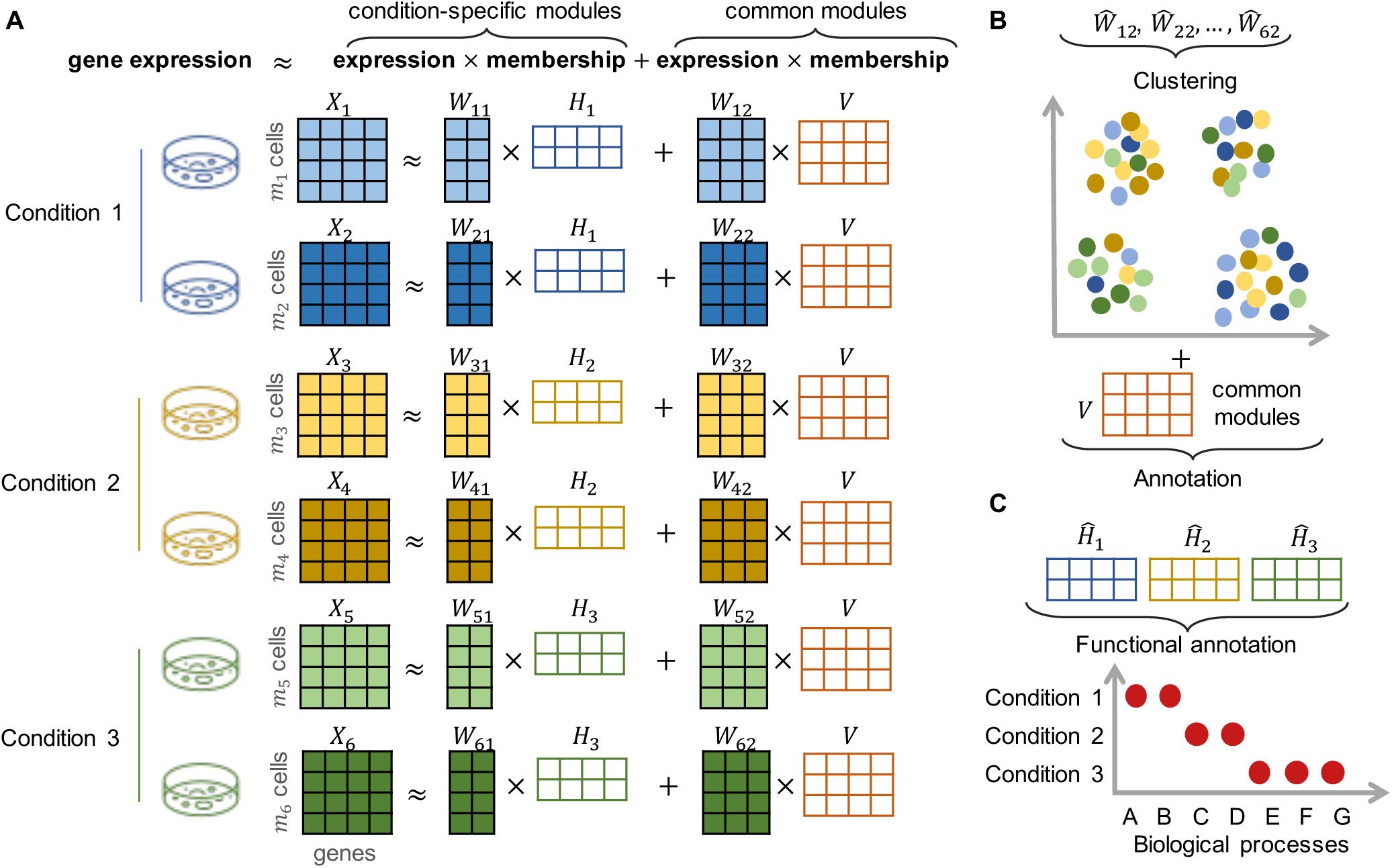
An overview of the scINSIGHT method. **A**: A toy example demonstrating the factorization model of scIN-SIGHT. In this example, we assume six single-cell samples from three biological conditions. Each sample, represented as a gene expression matrix (*X*_*ℓ*_, *ℓ* = 1, …, 6), is factorized into two components, the expression of two conditionspecific gene modules 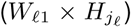 and the expression of three common gene modules (*W*_*ℓ*2_*× V*) (see Methods for details). **B**: After normalization (Methods), the inferred expression of common gene modules can be used to cluster cells and annotate cell types or states. **C**: The inferred condition-specific gene modules can be used to compare the transcriptome functions between conditions.

Our scINSIGHT method has the following features that distinguish it from existing integration tools developed for gene expression data. First, unlike existing NMF models such as iNMF [33], scCoGAPS [34], and SC-JNMF [35], scINSIGHT explicitly models both common and conditionspecific gene modules, allowing for the discovery of biologically meaningful differences among conditions and preventing them from being removed as technical effects (Methods). Second, unlike integration methods that construct normalized or integrated gene expression matrices in the original high-dimensional space [24, 25, 26], scINSIGHT achieves sparse, interpretable, and biologically meaningful decomposition of gene modules, which assist clustering and functional annotation. Third, the expression levels and memberships of common and condition-specific gene modules conveniently facilitate the comparison between samples and/or conditions in terms of cell cluster compositions and active biological processes.

### scINSIGHT reveals cellular identities by integrating simulated data across biological conditions

To benchmark the performance of scINSIGHT with ground truth information, we simulated synthetic single-cell gene expression data with known cell type compositions and condition-specific effects. Using our previously developed simulation tool scDesign [36], we simulated six single-cell samples from three time points (T1, T2, and T3), with two samples from each time point (see Methods for details). We considered six cell types, with three (C1, C2, and C3) present in all six samples and the other three (C4, C5, and C6) only present in particular conditions (Table 1). Before integration, the observed data presented distinct clusters corresponding to different cell types, samples, and conditions, making it difficult to identify genuine cell types across samples (Figure 2A).

**Table 1:** Cell type compositions in the six simulated single-cell samples.

| Condition | Sample | Cell type |  |  |  |  |  |
| --- | --- | --- | --- | --- | --- | --- | --- |
|  |  | C1 | C2 | C3 | C4 | C5 | C6 |
| T1 | S1 | 100 | 100 | 100 | 0 | 100 | 100 |
|  | S2 | 100 | 100 | 100 | 0 | 100 | 100 |
| T2 | S3 | 100 | 100 | 100 | 100 | 0 | 100 |
|  | S4 | 100 | 100 | 100 | 100 | 0 | 100 |
| T3 | S5 | 100 | 100 | 100 | 100 | 100 | 0 |
|  | S6 | 100 | 100 | 100 | 100 | 100 | 0 |

**Figure 2:**
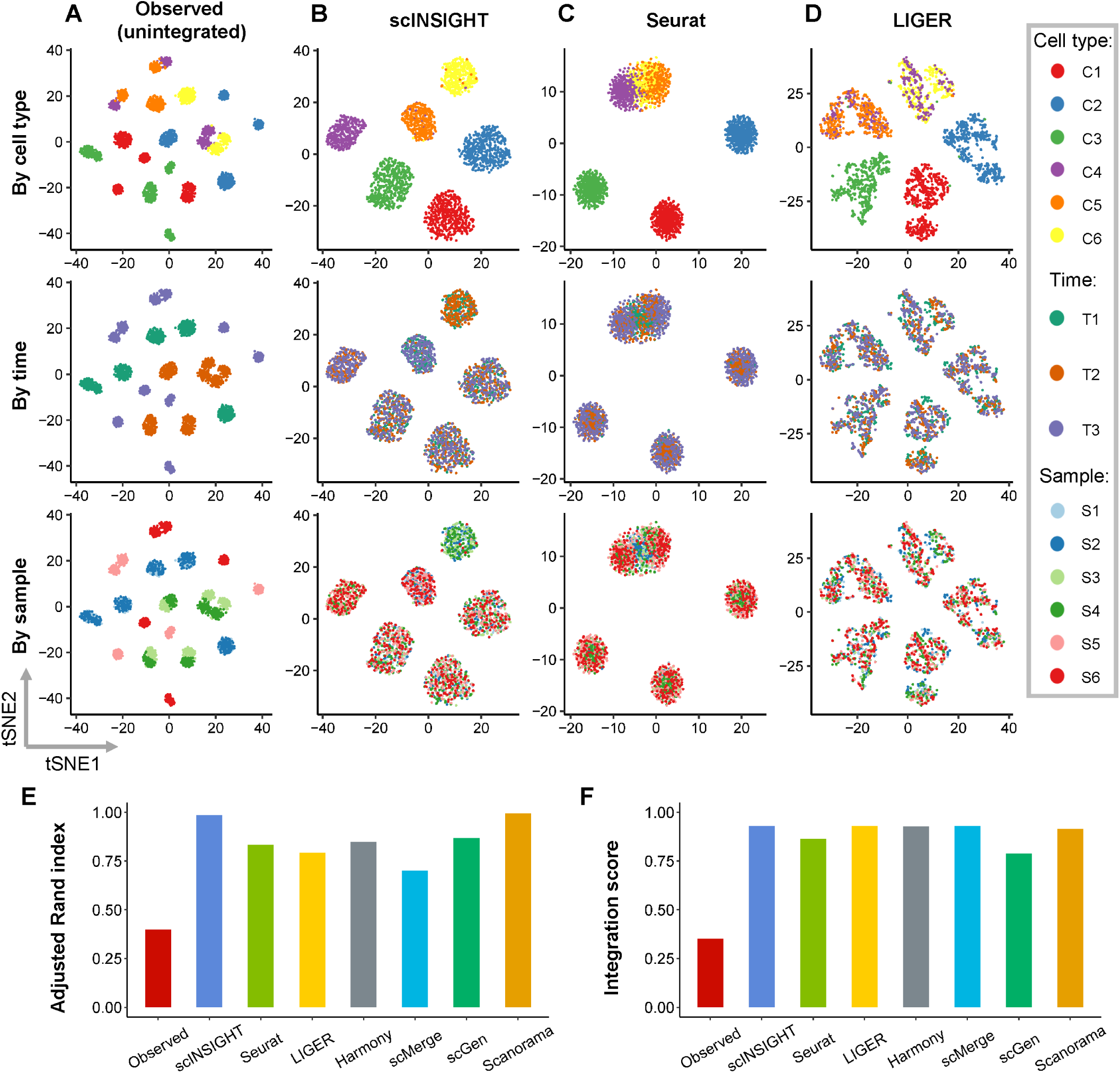
Comparison of observed and integrated data in the simulation study. **A-D**: tSNE plots of simulated cells based on the observed (unintegrated) data (**A**) and integrated data by scINSIGHT (**B**), Seurat (**C**), and LIGER (**D**). For each method, three tSNE plots colored by cell type, time point, or sample index are displayed. **E**: Adjusted Rand index calculated using clusters identified from the observed or integrated data. **F**: Integration scores of the observed and integrated data.

To obtain cell clusters that could represent real cellular identities, we applied scINSIGHT to the six gene expression samples, treating time point as the condition factor (Figure 2B, Methods). scINSIGHT identified six clusters based on the expression levels of 15 common gene modules, and the six clusters had a clear one-to-one correspondence with the ground truth cell types. For comparison, we also applied six alternative methods, Seurat [24], LIGER [27], Harmony [28], scMerge [26], scGen [37], and Scanorama [25] (Figures 2C-D and S1A-D), as they have shown preferable performance in a benchmark study of batch-effect correction methods for scRNA-seq data [30]. All methods were implemented following their suggested pipelines (Methods). As ground truth cell type labels were known, we calculated the adjusted Rand index (ARI) between inferred clusters and true labels, and scINSIGHT and Scanorama had the highest accuracy (0.99) in cluster detection (Figure 2E). To quantitatively compare the integration performance in terms of removing sample-specific technical effects, we defined an integration score to compare the frequency of cells from a sample in a local neighborhood with their frequency in the whole population (Methods). The integration score is between 0 and 1, and a score of 1 indicates full integration. The seven integration methods all demonstrated the ability to remove technical effects compared with the observed data (Figure 2F), and most methods had a score above 0.8. In addition, as the time-point effects on the gene expression mean were known in the simulation process, we compared the cosine similarity between the true time-point effects with the membership vectors of the inferred time-specific gene modules (Figure S1E), which confirmed that scINSIGHT is able to capture condition-specific gene modules.

To further evaluate the performance of scINSIGHT in different scenarios, we also considered three variants of the simulation study. Variant 1 represented the case when only one cell type (C1) was shared by all samples (C2 and C3 were removed). Variant 2 represented the case when there existed a rare common cell type (the cell number of cell type C2 was reduced to 20 in each sample). Variant 3 represented the case when there existed a rare condition-specific cell type (the cell number of cell type C4 was reduced to 20). Our results show that scINSIGHT achieved the second highest ARI after Scanorama in both variant 1 and variant 3 (Figures S2 and S3). In variant 2, scINSIGHTs clustering accuracy was similar to that of Seurat and LIGER and slightly lower than Harmony and Scanorama, but the common gene modules identified by scINSIGHT led to high-quality lower-dimensional visualization (Figure S4).

### scINSIGHT identifies T cell states associated with response to immunotherapy in melanoma

To assess the performance of scINSIGHT on real data, we first applied scINSIGHT to study immune cells in tumors of patients treated with checkpoint inhibitors such as anti-PD-1 and anti-CTLA4 [38]. In summary, we obtained single-cell gene expression data of 6,350 CD8+ T cells isolated from 48 tumor biopsies taken from 32 melanoma patients treated with the checkpoint therapy. Based on radiologic assessments, the 48 tumor samples were classified into 31 non-responders (NR) and 17 responders (R) [38]. The observed (unintegrated) data presented ten clusters (Methods), some of which were dominated by cells from a few donors (Figure 3A). For example, the cluster enclosed in the oval line mostly contained cells from a single donor, suggesting that the clustering analysis was affected by technical or donor-specific variability in the expression data.

**Figure 3:**
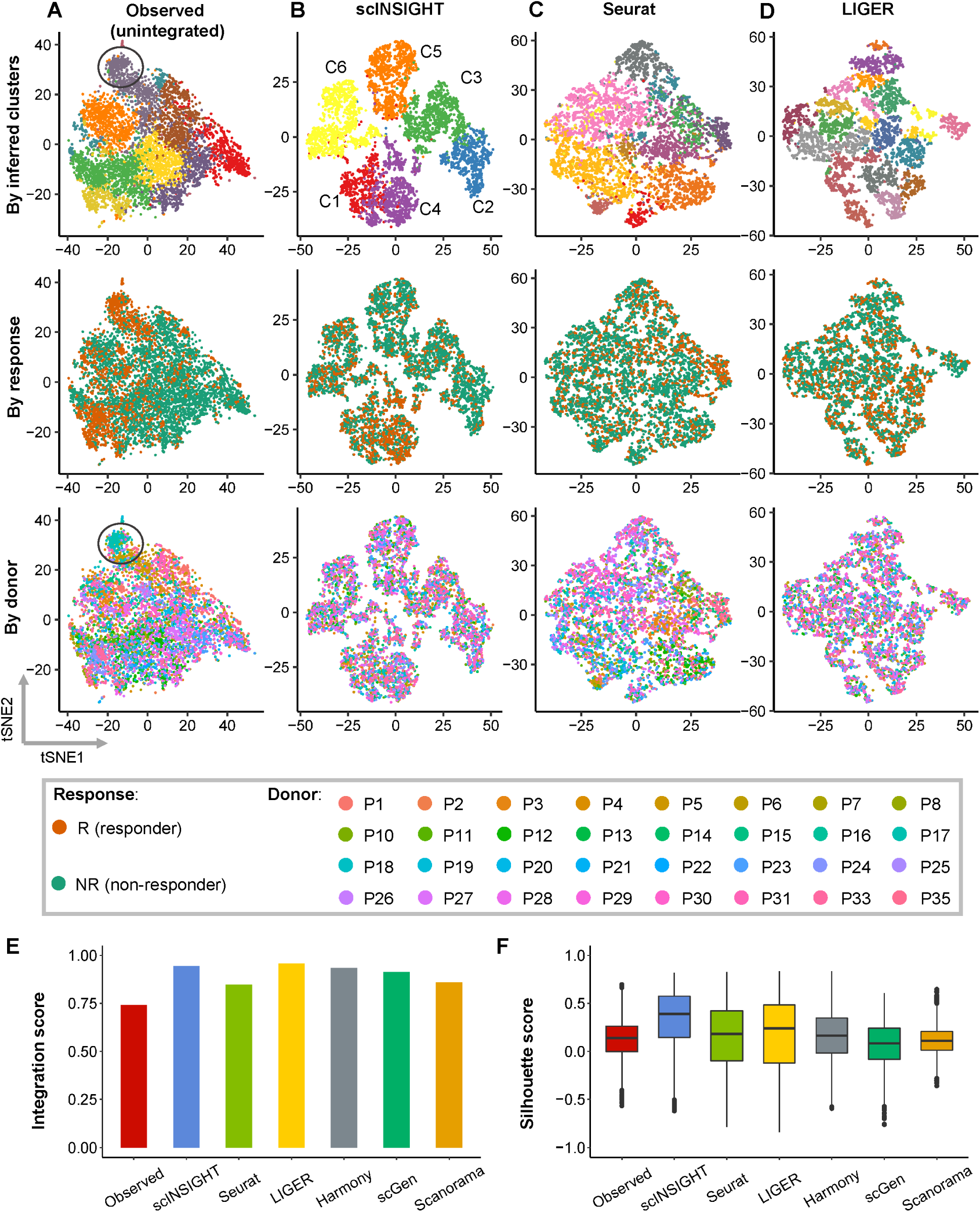
Comparison of observed and integrated data of CD8+ T cells from melanoma patients. **A-D**: tSNE plots of CD8+ T cells based on the observed (unintegrated) data (**A**) and integrated data by scINSIGHT (**B**), Seurat (**C**), and LIGER (**D**). For each method, three tSNE plots colored by inferred cell cluster, NR/R condition, or donor index are displayed. **E**: Integration scores of the observed and integrated data. **F**: Silhouette scores of the observed and integrated data.

To identify clusters corresponding to distinct T cell states and understand the biological difference between non-responder and responder samples, we applied scINSIGHT to the 48 single-cell samples (using expression of CD8+ gene signatures), treating NR/R as the condition factor (Methods). scINSIGHT identified six clusters (denoted as C1-C6) based on the activities of nine common gene modules (Figure 3B). Unlike the clusters in the unintegrated data, these six clusters do not represent obvious batch effect. We also applied Seurat, LIGER, Harmony, scGen, and Scanorama to the 48 samples using the same gene signatures (Methods; scMerge was not included since it encountered an error). These five methods identified 12, 19, 12, 11, and 11 clusters, respectively (Figures 3C-D and S5). To quantitatively assess the integration performance in terms of removing sample-specific technical effects, we calculated the integration score of each method. The six integration methods all demonstrated the ability to adjust for technical effects compared with the observed data (Figure 3E), and scINSIGHT (0.94) and LIGER (0.95) had the highest scores. To compare the consistency within the identified cell clusters, we calculated the Silhouette scores (Figure 3F), which suggested the highest consistency in clusters identified by scINSIGHT. In addition to using CD8+ gene signatures, we also repeated the above analysis using highly variable genes identified from the data (Methods). Compared with alternative methods, scINSIGHT still achieved the highest integration score and Silhouette scores (Figure S6).

Next, we investigated whether or not the computationally inferred clusters captured cell states associated with response to immunotherapy. For each cluster, we used a logistic regression model to evaluate if there was a significant association between the cluster proportion and the NR/R condition, while treating the donor information as covariates. Our analysis showed that four (66.7%) clusters identified by scINSIGHT were associated with response to immunotherapy (using a *P*-value threshold of 0.01), with two clusters (C1 and C4) enriched in responder samples and two (C2 and C5) enriched in non-responder samples (Figure 4A). The median percentage of C1 and C4 were 7.4% and 24.1% higher in responders, while the median percentage of C2 and C5 were 10.0% and 9.7% lower in responders, respectively. Among the 12 clusters identified by Harmony and 11 clusters by scGen, 41.7% and 45.5% clusters also had significantly different proportions between NR/R samples, respectively (Figures S7-S8). In contrast, the clusters identified from the integrated data by other methods either did not present association with response or had small difference between the NR/R samples (Figures S9-S11). For example, the difference in median percentage of the most significant cluster identified by Seurat, LIGER, and Scanorama were 1.7%, 5.6%, and 4.1%, respectively. This comparison suggests that scINSIGHT has improved sensitivity in detecting cell states enriched in specific biological conditions.

**Figure 4:**
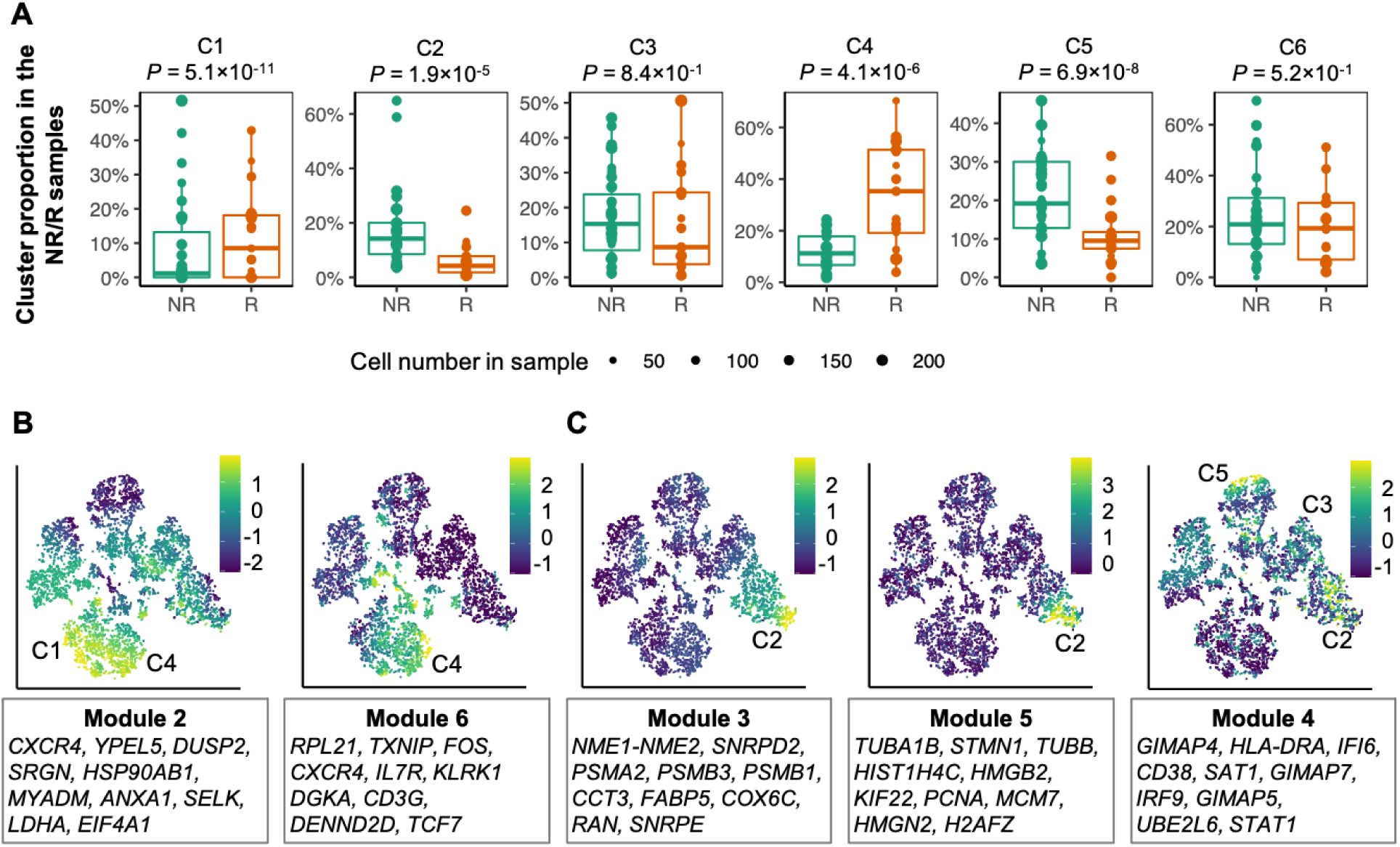
scINSIGHT identifies T cell states associated with immunotherapy response. **A**: Percentage of the six clusters identified by scINSIGHT in the 48 samples. *P*-values indicate significance of association between cluster proportion and immunotherapy response, and were calculated based on the logistic regression model. **B-C**: tSNE plots based on the integrated data by scINSIGHT. Color indicates scaled expression of the common gene modules.

We checked the nine common modules identified by scINSIGHT and found five modules highly expressed in the R-enriched or NR-enriched clusters. Modules 2 and 6 were highly expressed in C1 and C4 (enriched in responder samples) (Figure 4B). The ten genes with largest coefficients in these two modules contain seven and nine markers of natural killer T cells in the CellMarker database [40], respectively. In contrast, modules 3 and 5 were highly expressed in C2, and module 4 was highly expressed in C2, C3, and C5 (enriched in non-responder samples) (Figure 4C). In particular, the top ten genes in module 4 contain five markers of exhausted CD8+ T cells. We compared the scINSIGHT-inferred cell clusters with the cell states annotated in the original publication [38], and found that C3 and C5 corresponded to exhausted CD8+ T cells and C2 corresponded to exhausted lymphocytes (Figure S12). In contrast, C1 and C4 corresponded to lymphocytes and memory T cells. These findings are consistent with existing studies showing a correlation between T cell exhaustion and immune dysfunction in cancer [41]. We also compared the cell clusters identified by the observed data or the other methods with the cell type annotations, but they presented much lower consistency (Figures S12-S13). For the condition-specific gene modules, we compared the KEGG pathways [42] enriched in the top 100 genes with the largest coefficients in the R-specific or NR-specific gene modules identified by scINSIGHT. We found that PD-L1 expression and PD-1 checkpoint pathway in cancer and NF-kappa B signaling pathway were enriched in the R-specific module but not in the NR-specific module. In addition to the above results, we performed a co-expression analysis and confirmed that the identified gene modules presented stronger within-module co-expression than between-module co-expression (Figure S14).

### scINSIGHT identifies B cell types associated with disease phase of COVID-19 patients

To further evaluate the performance of scINSIGHT on complex data, we applied it to study B cells from peripheral blood samples of COVID-19 patients at different clinical phases [43]. We downloaded single-cell gene expression data of 9, 741 B cells from 14 blood samples of 13 donors. The 14 samples were divided into three phases: 5 healthy, 4 complicated (disease phase with severe signs of a systemic inflammatory response), and 5 recovery/pre-discharge (disease phase with no supplemental oxygen and absent inflammation markers) [43].

To investigate how B cell compositions and gene modules differ among the three phases, we applied scINSIGHT to the gene expression data of the 14 samples, treating disease phase as the condition factor (Methods). scINSIGHT identified 13 common gene modules and six phase-specific gene modules, which also demonstrated stronger within-module co-expression than between-module co-expression (Figure S15). Based on the common gene modules, scINSIGHT discovered ten B cell clusters across the three phases (Figure 5A). To annotate the B cell clusters with major B cell types, we used the SingleR [44] method to classify the cells by comparing their transcriptomes with bulk data references. We found a clear correspondence between scINSIGHT’s cluster assignments and reference-based annotations, with C3 matched with naive B cells, C5 and C10 matched with plasma B cells, and the other clusters matched with memory B cells (Figure 5A). For comparison, we also performed the analysis using the observed data or integrated data by the six alternative methods (Figures 5B and S16). We calculated the ARI between the cluster assignments of each method and the reference annotations, and scINSIGHT led to the best consistency (Figure 5C). As the clusters identified from single-cell data might correspond to B cell subtypes not available in the bulk reference, it was expected that the overall ARI was not very high. We also compared the integration score of the seven methods (Figure 5D), and all showed better performance than directly using the observed data.

**Figure 5:**
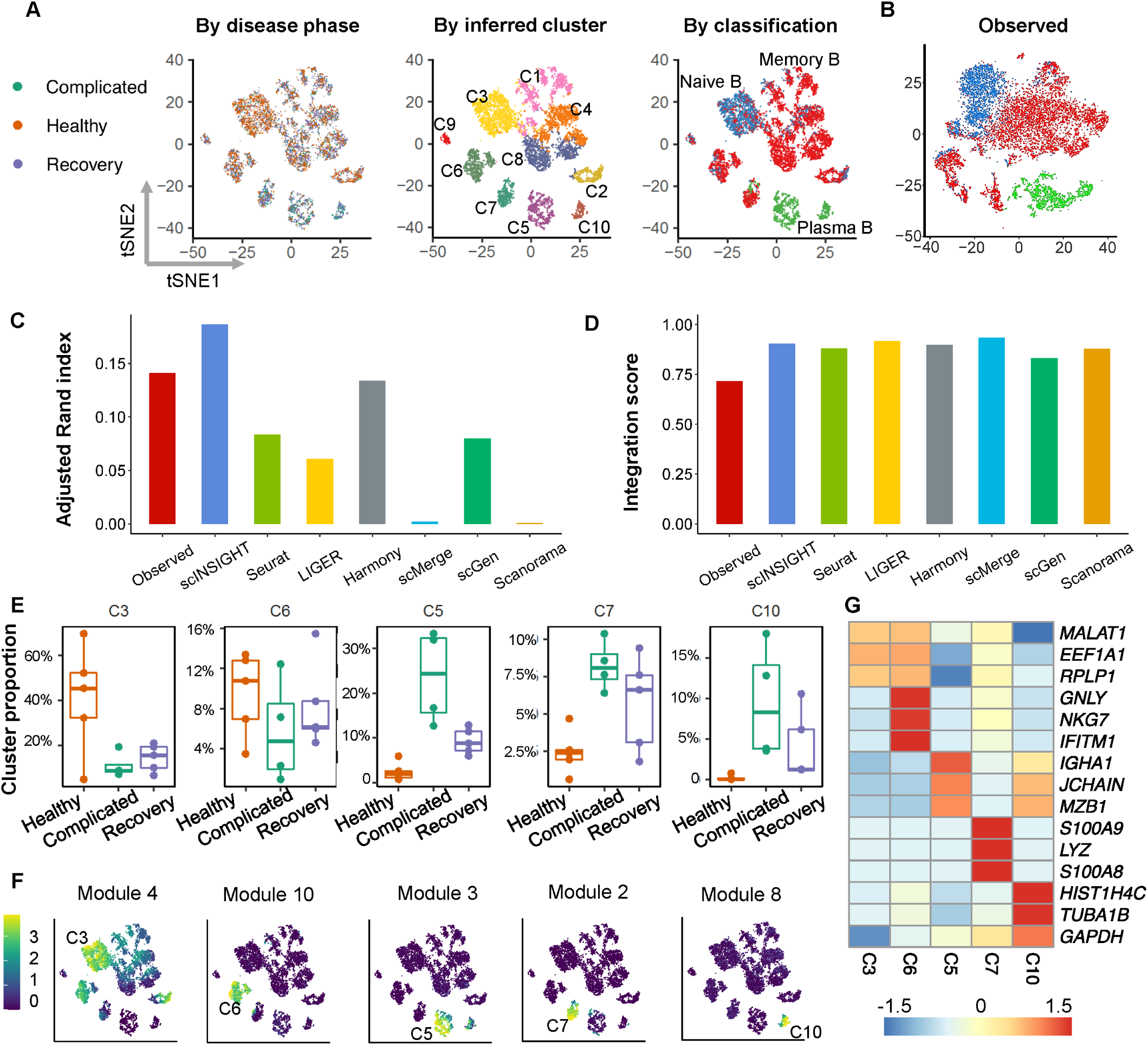
Comparison of observed and integrated data of B cells from COVID-19 patients. **A**: tSNE plots of B cells based on integrated data by scINSIGHT. Cells are colored by disease phase, inferred cluster membership, or classified cell type (by SingleR). **B**: tSNE plots of B cells based on the observed (unintegrated) data. Cells are colored by classified cell type (by SingleR). **C**: Adjusted Rand index of inferred clusters compared with the SingleR classification. **D**: Integration scores of the observed and integrated data. **E**: Percentage of five clusters identified by scINSIGHT in the 14 samples. **F**: Expression levels of five common modules that were specifically highly expressed in the five clusters in panel **E. G**: Expression of the top three genes with the largest coefficients in modules 4, 10, 3, 2, and 8. Expression levels were averaged within each cluster and then scaled across clusters.

Next, we compared the proportion of the ten scINSIGHT-inferred clusters among the three phases. For each cluster, we used a logistic regression model to evaluate if there was a significant association between the cluster proportion and the clinical phase, after accounting for the donor factor. We found two clusters, C3 (naive B) and C6 (memory B), enriched in healthy samples, and three clusters, C5 (plasma B), C7 (memory B), and C10 (plasma B), enriched in complicated samples (Figure 5E). This is consistent with recent studies showing that protective immunity induced by COVID-19 infection may rely on the production of both memory B cells and plasma cells [45, 46]. In addition, we observed that the median proportion of these B cell clusters in recovery/pre-discharge samples was always between their median proportions in healthy and complicated samples, suggesting that gene expression profiles during recovery from COVID-19 carry characteristics of both healthy and complicated clinical phases.

To understand the transcriptome difference between the above five B cell clusters, we identified representative gene modules for each cluster based on the expression of the 13 common modules detected by scINSIGHT (Figure 5F). The results confirmed that C6 and C7 were characterized by distinct modules and genes (Figure 5G). Module 10 was specifically highly expressed in C6, and had enriched GO terms associated with activating signal transduction and T cell activation; module 2 was specifically highly expressed in C7, and had enriched GO terms associated with regulation of inflammatory response and neutrophil chemotaxis (Figure S17). As both *GNLY* and *NKG7* had relative high expression in C6, we hypothesize that C6 represented a natural-Killer-like B cell population [47], or it was annotated as B cell by mistake. In addition, C5 and C10 were two subtypes of plasma cells expressing modules with distinct functions. C5 was characterized by module 3, which was associated with endoplasmic reticulum to cytosol transport, while C10 was characterized by module 8, which was related to ATP synthesis and oxidative phophorylation (Figure S17).

We also compared the expression of phase-specific modules identified by scINSIGHT, and confirmed that genes with large coefficients in a condition-specific module indeed had higher expression levels in samples belonging to that condition (Figure 6A). To compare the biological functions of the healthy-specific, complicated-specific, and recovery-specific modules, we performed pathway enrichment analysis using the 100 genes with the largest coefficients in each module. We found that the healthy-specific and complicated-specific modules had distinct sets of enriched terms, and the recovery-specific module had overlapping terms with the former two in addition to its unique terms (Figure 6B). In particular, the complicated-specific module was enriched with genes involved in interferon signaling, antigen presentation, and ATF6 activation, which play key roles in innate immune response; the recovery-specific module was enriched with genes involved in ER-to-Golgi transport and cell cycle, which is consistent with the observations in an independent COVID-19 study [50]. The above results demonstrate scINSIGHT’s ability to help compare active biological processes between biological conditions, which cannot be achieved by analyzing individual samples or by existing integration methods.

**Figure 6:**
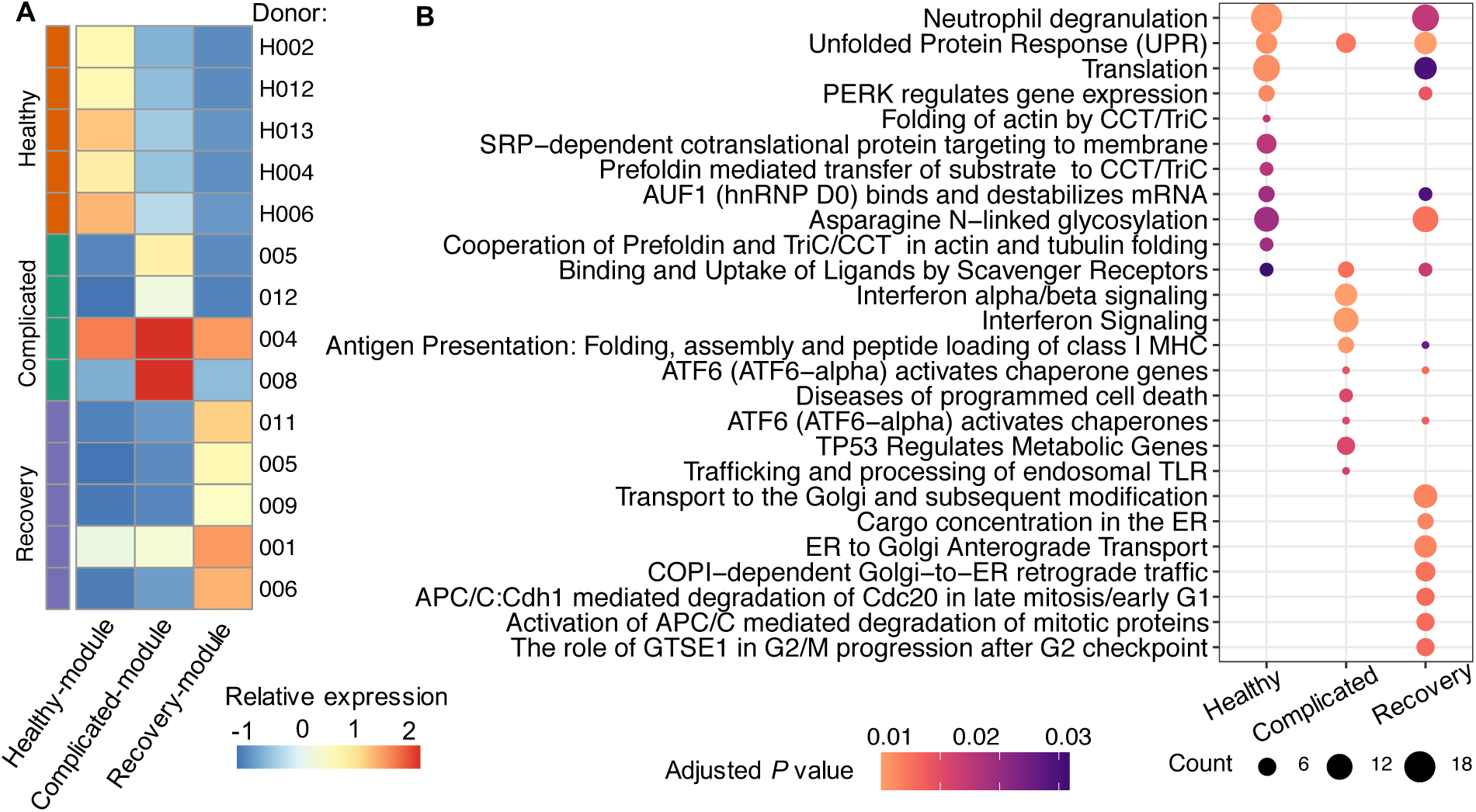
Comparison of disease phase-specific modules identified by scINSIGHT. **A**: Expression levels of condition-specific module identified by scINSIGHT. For each module, we calculated the expression of the 100 genes with the largest coefficients based on *Ŵ* _*ℓ*1_*Ĥ*_*ℓ*_ (*ℓ* = 1, …, 14), and averaged the expression across single cells in each sample. **B**: Top 10 enriched REACTOME pathways [48, 49] in the condition-specific modules identified by scINSIGHT.

### scINSIGHT detects dermal cell populations during murine skin wound healing

Lastly, we assessed the performance of scINSIGHT in a special case where only a single sample is available from each condition. In this application, the condition-specific components might capture both biological and technical differences in single-cell transcriptomes, so we focused on the common gene modules identified by scINSIGHT when investigating the results. The gene expression dataset we used was sequenced from dermal cells of wound dermis from mice in control or Hedgehog (Hh) activation conditions [51]. The activation of the Hh pathway has been shown to induce hair follicle regeneration during murine skin wound healing [51].

We applied scINSIGHT to the gene expression data of the two samples, treating control/treatment as the condition factor (Methods). scINSIGHT identified 11 common gene modules and four condition-specific gene modules. As in the previous applications, we observed stronger within-module co-expression than between-module co-expression (Figure S18). Based on 11 common gene modules, scINSIGHT discovered 13 cell clusters across the two conditions. Using the average expression levels of lineage-specific gene signatures [51], we annotated the cell clusters as six cell types (Figure 7A): Hh-activated fibrolasts (*Ptch1, Lox, Dpt*), Hh-inactive fibrolasts (*Lox, Dpt*), muscle cells (*Myh11, Rgs5, Notch3*), endothelial cells (*Pecam1, Cdh5, Vwf*), Schwann cells (*Plp1, Mbp, Sox10*), and immune cells (*Cd68, H2-Aa*). We also performed the analysis on the observed data and integrated data by Seurat, LIGER, Harmony, scMerge, scGen, and Scanorama (Figure S19). scINSIGHT achieved the highest Silhouette scores among all the methods, and the second highest integration score after scMerge (Figure 7B-C). Nevertheless, the low Silhouette scores of scMerge suggest that it removed biologically meaningful signals during integration. We compared the inferred expression of the 11 common gene modules and found that module 1 was highly expressed in Hh-inactive fibroblasts and modules 5 and 11 were highly expressed in Hh-active fibroblasts (Figure 7D). The comparison also indicated the existence of two sub-populations in Hh-active fibroblasts, with differential expression of modules 5 and 11. GO enrichment analysis showed that module 1 was relevant to extracellular matrix organization and collagen-related biological processes, which are fundamental to fibroblast proliferation (Figure 7E). In contrast, module 5, highly expressed in one sub-population of Hh-active fibroblasts, was associated with epithelial cell proliferation and cellular responses to metal ions, which promote wound healing. Module 11 was highly expressed in the other sub-population of Hh-active fibroblasts, and it demonstrated overlapping biological functions with both modules 1 and 5. The results of scINSIGHT together revealed transition of cell fate in fibroblasts induced by Hh-activation, by integrating data across the two conditions and identifying common gene modules. In addition, we found that the clusters inferred by scINSIGHT demonstrated more specific expression of the fibroblast signatures *Lox* and *Dpt* and the Hh-active fibrolast signature *Ptch1* (Figure S20), suggesting that scINSIGHT had a high accuracy in detecting cell populations across biological conditions.

**Figure 7:**
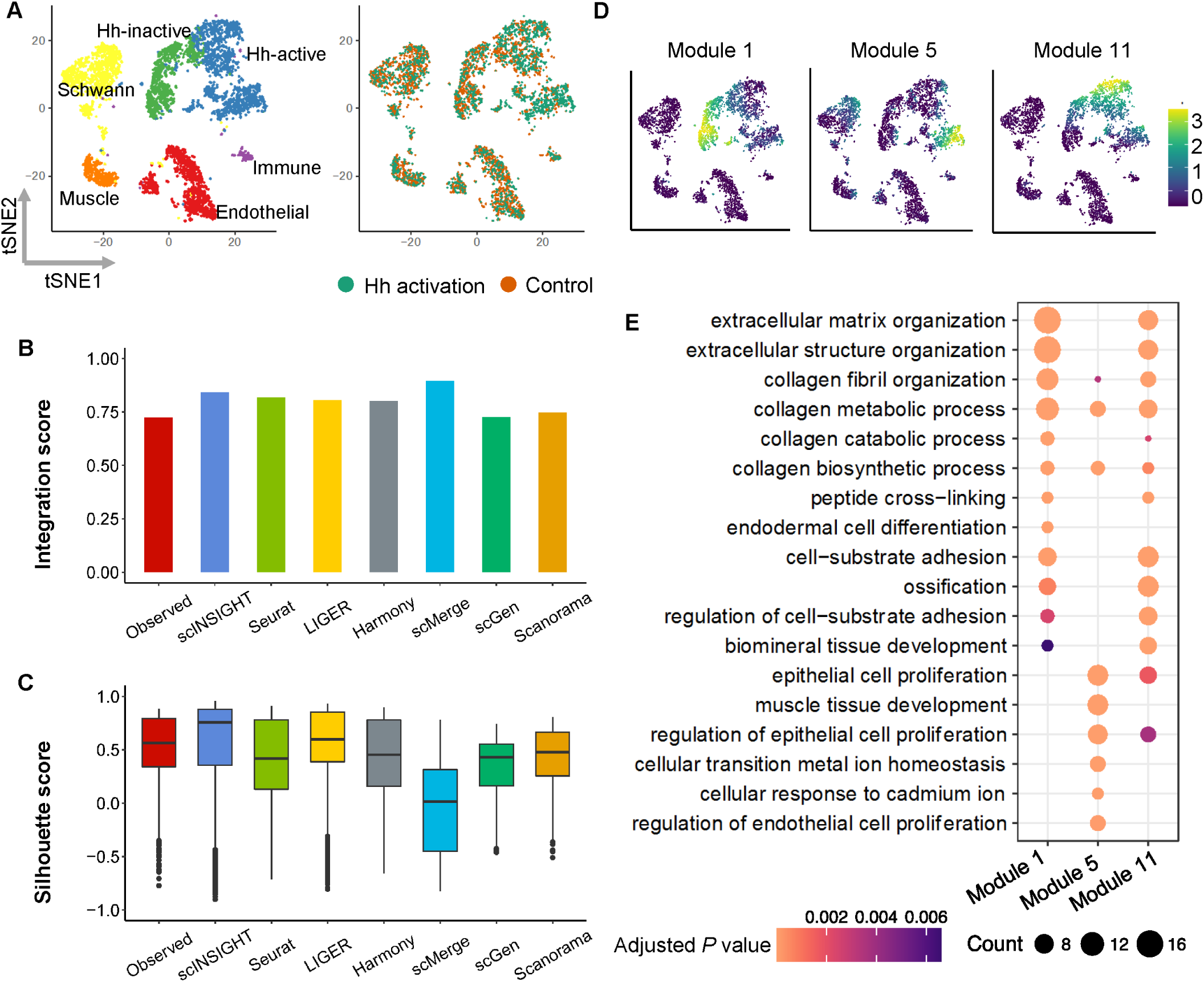
Comparison of observed and integrated data of dermal cells. **A**: tSNE plots of dermal cells based on integrated data by scINSIGHT. Cells are colored by cell type or activation condition. **B**: Integration scores of observed and integrated data. **C**: Silhouette scores of observed and integrated data. **D**: Expression levels of three common modules that were specifically highly expressed fibroblasts. **E**: Top enriched GO terms in modules 1, 5, and 11.

### Computational time and memory usage

We recorded the running time and memory usage of scINSIGHT and the other six methods (Seurat, LIGER, Harmony, scMerge, scGen and Scanorama) in Table S1. In addition to datasets discussed in previous applications, we also considered a mouse retina dataset with two samples [52, 53], 12 cell types, and 71,638 cells (Figure S21; Methods). Among the seven methods, scINSIGHTs memory usage is on the medium level. Regarding the running time, admittedly, scINSIGHT is slower than the other methods, and we summarize the major reasons below. First, it requires a more complex model to distinguish between technical signals, common biological signals, and condition-specific signals. Compared with LIGER which is also an NMF-based method, scINSIGHTs optimization is naturally more complex and takes more time. Second, scINSIGHT includes internal steps to identify optimal parameters, so its running time is expectedly larger than methods using given parameters. For example, LIGER has a function to select the dimensionality of factorized matrices, but this function encountered errors for multiple datasets, and we had to directly input the parameters in our analysis (Methods). scMerge also requires users to input the cluster number in each single-cell sample. Even though scINSIGHT takes relatively more time than the other methods, its performance is still acceptable to server users. Its overall running time for datasets with 3,000 cells (6 samples), 4,680 cells (2 samples), 6,350 cells (48 samples), 9,741 cells (13 samples), and 71,638 cells (2 samples) are 4.5 h, 0.9 h, 2.8 h, 2.9 h, 36.4 h, respectively. To further reduce the computational time of scINSIGHT given large-scale data, we have two suggestions for users. First, users can assign single cells to major cell types with well-known marker genes, and then apply scINSIGHT to single-cell samples of the same major cell type to reveal finer distinctions between cellular identities that cannot be determined using existing marker genes. Second, users can apply the Metacell [54] method separately to each single-cell sample to group statistically equivalent cells into metacells, thus reducing cell number in each sample, and then apply scINSIGHT to samples of metacells.

## Discussion

In this article, we propose a new method named scINSIGHT to address the problem of integrating heterogeneous single-cell data from multiple biological conditions (i.e, groups). Based on a novel matrix factorization model developed for analyzing multiple single-cell samples, scINSIGHT learns coordinated gene expression patterns that are common among or specific to different biological conditions, offering a unique chance to jointly identify heterogeneous biological processes and diverse cell types. In addition, its identified gene expression patterns provide a convenient way to compare single-cell samples across conditions such as different experimental groups, time points, or clinical conditions.

We benchmarked the performance of scINSIGHT in both simulation and real data studies, in comparison with analysis without integration or with six alternative integration methods. Using the ground truth information in simulation as a reference, we confirmed scINSIGHT’s ability to accurately decompose common and condition-specific gene modules, and to precisely identify cellular identities based on the inferred expression of common gene modules. In the three real data applications, scINSIGHT repeatedly demonstrated its effectiveness to analyze, compare, and interpret single-cell gene expression data across samples and biological conditions. Based on its identified cell clusters and decomposed gene modules, scINSIGHT is able to discover T cell states associated with response to immunotherapy in melanoma patients, B cell types associated with disease phase of COVID-19 patients, and dermal cell populations for murine skin wound healing. In addition, scINSIGHT consistently showed higher accuracy and interpretability than the other methods in the above real data studies.

Based on the application to both simulated and real data, we summarize the advantages of scINSIGHT as follows. First, it jointly defines cellular identities across multiple single-cell samples, accompanied by characteristic gene modules which enable straightforward and transparent interpretation of each cell cluster’s function. Second, as cellular identities are inferred based on common gene modules and not biased by sample-specific or condition-specific effects, scINSIGHT allows accurate comparison of cell composition across samples and biological conditions. Third, the condition-specific gene modules provide biological insights towards gene expression mechanisms in distinct but related conditions, after adjusting for the difference in cell composition. The above information is challenging to obtain if the single-cell samples are analyzed individually. A future direction to further improve the flexibility of scINSIGHT is to adjust its regularization term in model (2) to account for more complex relationships among the biological conditions. For example, when single-cell samples are from multiple clinical phenotypes with a hierarchical structure of an ontology, such information can be incorporated into the regularization term to more accurately identify condition-associated gene modules.

We would like to point out that scINSIGHT is not intended to replace existing batch-effect removal tools. If it is known that the single-cell samples are from the same tissue types, biopsies, or cell lines, and the main difference lies in the sequencing platform or library preparation protocol, then it would suffice to use existing batch-effect removal or integration methods to remove unwanted technical variation from the gene expression data. Additionally, if there are known biological differences which are not of interest, existing methods might also achieve desirable results to align single-cell samples. Yet in this case scINSIGHT is expected to identify cell populations with higher accuracy, as clustering is performed after condition-specific effect is identified and removed.

In addition to single-cell gene expression data, several integration methods have been used to integrate multi-omics single-cell data, including those of DNA methylation, chromatin accessibility, *in situ* gene expression, and protein expression [24, 27, 55]. A potential extension of the scINSIGHT model is to use common and modality-specific factors to respectively account for highly correlated and uncorrelated expression patterns in data samples from different assays. With this extension, we might use scINSIGHT to define cellular identities with multi-omics data and to understand regulatory mechanisms of gene expression and/or protein synthesis.

## Methods

### The scINSIGHT model

We propose a matrix factorization model named scINSIGHT to jointly analyze multiple single-cell gene expression samples from different biological conditions (i.e., groups). These biological conditions can be assigned based on phenotypes or experimental conditions, such as different developmental stages, disease phases or treatment groups. It aims to simultaneously identify common and condition-specific gene modules and quantify their expression levels in each sample in a lower-dimensional space. Each gene module is represented as a non-negative linear combination of a subset of coordinated genes. The common gene modules are universally expressed across conditions, while the condition-specific gene modules are only highly expressed in specific conditions.

We first introduce the scINSIGHT model and algorithm, then provide more details of data processing and parameter selection in the subsequent sections. Suppose there are *L* single-cell gene expression samples obtained from biologically heterogeneous sources. Each sample is represented by a gene expression matrix, after read mapping and proper normalization. We assume that these *L* samples can be divided into *J* biological conditions (*J* ≤ *L*). For example, the biological conditions may correspond to experimental groups (e.g., case and control), time points, or donor categories. We use index *j*_*ℓ*_ ∈ 1, 2, …, *J* to denote the condition which sample *ℓ* belongs to. Without loss of generality, we assume the columns in these matrices represent genes and the rows represent individual cells. We further assume that the *L* matrices share the same set of genes which are listed in the same order. Then, our scINSIGHT model specifies that, for sample *ℓ* (*ℓ* = 1, 2, …, *L*),

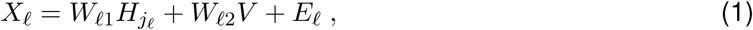

where 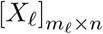 is the gene expression matrix with *m*_*ℓ*_ cells and *n* genes for sample *ℓ*;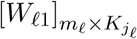 is the expression matrix of 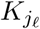 condition-specific gene modules for sample 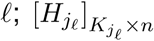 is the membership matrix of 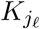 condition-specific gene modules for condition *j*, and it’s shared by all samples belonging to condition 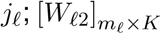 is the expression matrix of *K* common gene modules for sample *ℓ*; [*V*]_*K×n*_ is the membership matrix of *K* common gene modules, and it’s shared by all *L* samples; *E*_*ℓ*_ is the residual matrix for sample *ℓ*. In addition, we require all the matrices to be non-negative. The above model allows scINSIGHT to detect common gene modules by borrowing information across samples and condition-specific gene modules by borrowing information among samples of the same condition.

In order to solve the scINSIGHT model and identify the common and condition-specific modules, we formulate a minimization problem with an objective function in the following form

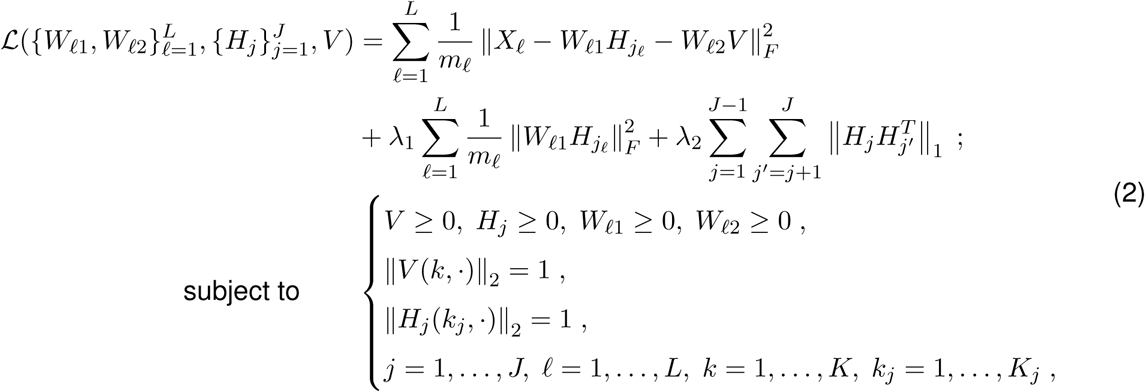

where *A*(*k*,) denotes the *k*-th row of matrix *A*, ∥·∥_*F*_ represents the Frobenius norm, ∥·∥_1_ represents the sum of all elements of a matrix, and ∥·∥_2_ represents the *L*_2_ norm of a vector. The first term in the objection function aims to minimize the differences between observed and reconstructed gene expression matrices; the second term serves as a regularization term to control the scale of condition-specific components. In the first and second terms, the Frobenius norms are scaled by cell numbers in each sample to prevent the results from being dominated by large samples.

To obtain the optimal solutions for problem (2), we have developed the following algorithm based on the block coordinate descent framework [32]. We present the major update steps below, and the detailed algorithm and derivation are introduced in the Supplementary Methods.

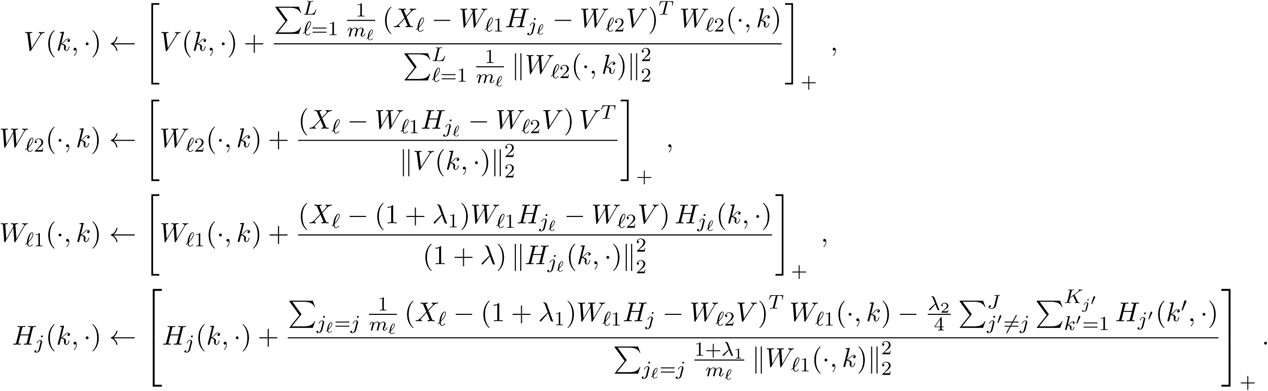

For comparison purpose, we also introduce the formulation of the iNMF model [33] used by the LIGER method. It assumes the following relationship for sample *ℓ* (*ℓ* = 1, 2, …, *L*),

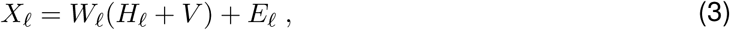

where 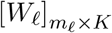 is the expression matrix for gene modules in sample 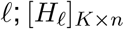 is the membership matrix for dataset-specific gene modules in sample *ℓ*;[*V*]_*K*×*n*_ is the membership matrix for shared gene modules. To solve the model, the objective function is formulated as:

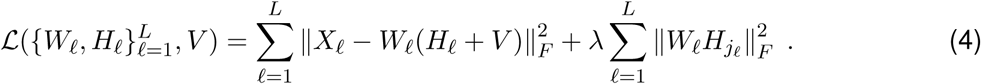

Unlike the scINISGHT model, model (3) has the following limitations: (1) it does not account for the assignment of biological conditions; (2) it assumes that there exists the same number of shared gene modules and dataset-specific gene modules; (3) it cannot decompose the expression of shared and dataset-specific gene modules. These limitations are resolved by scINSIGHT to better jointly analyze single-cell samples from different biological conditions.

### Processing of read count matrices

Given the *L* read or UMI count matrices, we use the following three steps to obtain the *L* processed gene expression matrices. First, the count matrices are normalized by cell library sizes. We scale the counts such that each cell has a total of 10^5^ reads or UMIs. Second, we perform log-transformation on the scaled values. Third, we remove genes that exhibit low variation among individual cells to increase the signal-to-noise ratio in gene expression data. The highly variable genes are selected using the same approach as described in Seurat [56]. If users prefer a different normalization or gene selection procedure [57], the scINSIGHT software also allows users to supply normalized and filtered gene expression matrices.

### Cell clustering based on common gene modules

As the NMF framework has an inherent clustering property, we can use the expression of common gene modules to cluster single cells from different samples, and use the membership of common gene modules to interpret cellular identities. However, since minimizing the objective function in problem (2) does not directly ensure an optimal clustering performance, we propose a cell clustering method based on 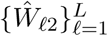. The method was inspired by previous batch correction methods [18, 27], and it performs alignment using mutual nearest neighbors in the space of 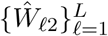. It has the following key steps.

1. The row vectors in 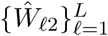 are normalized such that their *L*_2_ norms all equal 1. With a slight abuse of notation, we still use 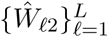 to denote the expression of common gene modules after normalization.
2. Given every pair of samples *ℓ* and *ℓ*, (*ℓ, ℓ*, ∈ {1, …, *L*}), from sample *ℓ*, we find the *n*_*c*_ nearest neighbors for each cell in sample *ℓ*. When searching for nearest neighbors, the cell-cell distance is calculated using the Euclidean distance between normalized module expression in *Ŵ* _*ℓ*_ and *Ŵ* _*ℓ*_ ^*′*^ 2. By default, *n*_*c*_ = 20.
3. A mutual nearest neighbor graph is constructed using cells in all *L* samples as the nodes. Cell *i* in sample *ℓ* and cell *i*, in sample *ℓ*, are connected in the graph if and only if they are mutual nearest neighbors.
4. The Louvain method [58] is used to perform clustering on the mutual nearest neighbor graph. The clustering result is then used to label cells across samples.
5. To produce visualization that better reflects the clustering result, we align the module expression within each cell cluster using quantile normalization. We denote the quantile normalized module expression as 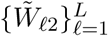, and they are used for downstream visualization.

### Selection of model parameters

We propose a heuristic method to first select *K*, the number of common gene modules, and then select *λ*_1_ and *λ*_2_, the two regularization parameters in the scINSIGHT model (2). The numbers of condition-specific gene modules, 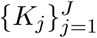, are expected to be small compared with *K*. We set *K*_*j*_ = 2 (*j* = 1, …, *J*) in our analysis to facilitate interpretation of factorization results. To select *K* among a set of candidate values, we calculate a stability score inspired by consensus clustering [59] and choose the value that leads to the largest stability score. For every candidate value of *K*, we run our algorithm *B* (defaults to 5) times with different initializations and *λ*_1_ = *λ*_2_ = 0.01.1, …, *B*), We represent the clustering result from each run as a binary matrix 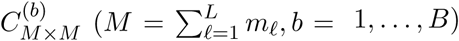, with 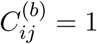 indicating that cells *i* and *j* belong to the same cluster. We then calculate a consensus matrix 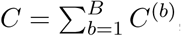, whose entries range between 0 and 1 and reflect the probability that two cells belong to the same cluster. The stability score is defined as the Pearson correlation between corresponding entries in two matrices, *I*_*M×M*_ − *C* and the cophenetic distance matrix (for hierarchical clustering on *C*) [60]. A large (close to 1) stability score indicates that the assignment of cells to clusters varies little with different initializations, suggesting a strong clustering pattern with the corresponding *K* value. Therefore, we select the value of *K* that leads to the largest stability score.

Next, we select the regularization parameters *λ*_1_ and *λ*_2_, fixing *K* to the selected value. In our simulation study, we found that *λ*_1_ has a much greater effect on scINSIGHT’s results than *λ*_2_, so we set *λ*_1_ = *λ*_2_ to simplify the computation. We consider five candidate values, {0.001, 0.01, 0.1, 1, 10}, and select the optimal value based on a specificity score. The specificity score indicates how well the condition-specific gene modules capture specific gene expression patterns in each condition. For the *k*-th gene module in *Ĥj*, we find the 100 genes that have the largest coefficients on the module, and use 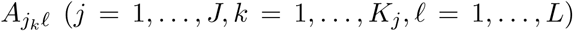 to denote the average of their expression across all cells in each sample. The specificity score is then defined as

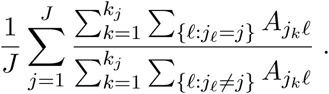

We select the regularization value that leads to the largest specificity score.

### Calculation of evaluation metrics

The integration score aims to quantify how well the (unintegrated or integrated) data removes sample-specific technical effects. We consider cells from all samples, and use *ℓ*_*i*_ to denote the sample index of cell 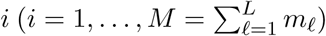. In addition, *N*_*i*_ represents the *k*-nearest neighbor set of cell *i*. For observed (unintegrated) data, *N*_*i*_ is defined based on cell-cell distance calculated using the first 20 PCs. For scINSIGHT, *N*_*i*_ is based on cellular distances calculated using 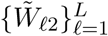.

For Seurat and LIGER, *N*_*i*_ is also determined based on the gene expression data after integration. For each set of *N*_*i*_’s, the corresponding integration score is defined as

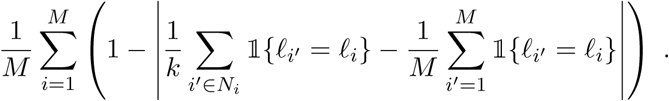

The integration score is among 0 and 1, and it compares the frequency of cells from a sample in a local neighborhood with their frequency in the whole population. An integration score of 1 indicates full integration.

The Silhouette score was calculated using the R package `cluster` and the adjusted Rand index was calculated using the R package `mclust`.

### Simulation and analysis of synthetic data

To better benchmark the performance of scINSIGHT, we used the scDesign method [36] to simulate high-quality synthetic single-cell gene expression datasets that capture the distribution characteristics of real data. In summary, we simulated six single-cell samples from three time points (T1, T2, and T3 corresponding to three biological conditions), with two samples from each time point. To reflect different cell type compositions among the time points, we assumed that three cell types (C1, C2, and C3) were present in all three time points, while samples in T1 didn’t have cell type C4, T2 didn’t have C5, and T3 didn’t have C6.

As scDesign learns gene expression parameters from real data to simulate new synthetic cells, we used the following approach to incorporate biological heterogeneity into the simulated data. First, the expression parameters corresponding to the cell type effect were learned from six immune cell types in a dataset of peripheral blood mononuclear cells [61]. Second, the expression parameters corresponding to the time point effect were learned from a time-course scRNA-seq dataset [62]. Third, for each synthetic cell, its mean gene expression was calculated as a weighted average of the corresponding cell type mean effect (weight of 0.9) and the time point mean effect (weight of 0.1) estimated by scDesign. Fourth, we used scDesign to generate simulated gene expression matrices for the six samples based on the expression mean and other learned expression parameters.

For analysis based on the observed (unintegrated) data, we calculated the first 20 principal components (PCs) after library size normalization and log-transformation, and used the cells’ scores on the PCs to calculate tSNE coordinates and perform Louvain clustering [58]. scINSIGHT was applied to the six samples treating time point as the condition factor. We set *K*_*j*_ = 2 for all cases. The best *K* was 9 for variant 2 and 15 for the other cases (selected from {5, 7, 9, 11, 13, 15}). For all cases, the regularization values were selected as *λ*_1_ = *λ*_2_ = 0.01. LIGER was applied with nrep=5 and k=11, the other parameters were set as the default values. Seurat, Harmony and Scanorama were applied with default parameters. scMerge was applied with kmeansK=rep(3,6) in variant 1 and kmeansK=rep(5,6) in all other cases. The other parameters were set as the default values. Follow its tutorial, scGen was applied with max epochs=100 and batch size=32, early stopping=True, and early stopping patience=25, and the other parameters were set as the default values. For clustering, scINSIGHT, Seurat, and LIGER have their own clustering methods, Harmony used Seurat’s clustering in its tutorial, and Scanorama used the Leiden algorithm [63] in its tutorial. For these methods, we used their default methods or methods demonstrated in their tutorials to perform clustering. Since scMerge and scGen do not provide clustering functions, we applied the Louvain algorithm [58] to perform clustering on their integrated data..

The scINSIGHT, LIGER, Seurat, Harmony, scMerge, scGen, and Scanorama methods were applied using R package `scINSIGHT` (v0.1.1), R package `rliger` (v0.5.0), R package `Seurat` (v3.2.3), R package `harmony` (v0.1.0), R package `scMerge` (v1.6.0), Python package `scgen` (v2.0.0), and Python package `Scanorama` (v1.7.1), respectively.

### Analysis of real data

For the melanoma dataset [38], we processed the data as described above, treating each biopsy as one sample. For analysis with CD8+ gene signatures, the gene features were obtained from Sade-Feldman et al [38]. The normalized expression data with selected gene features were used for data integration with all methods. For analysis based on the observed (unintegrated) data, we calculated the first 20 PCs and used the cells’ scores on the PCs to calculate tSNE coordinates and perform Louvain clustering. For scINSIGHT analysis, scINSIGHT was applied to the 48 samples treating response as the condition factor. We set *K*_*j*_ = 2 and the best *K* was 9 (selected from {5, 7, 9, 11, 13, 15}). The regularization values were selected as *λ*_1_ = *λ*_2_ = 1. LIGER was applied with nrep=5 and k=5 as larger values would lead to errors, the other parameters were set as the default values. Seurat, Harmony, and Scanorama were applied with default parameters. scGen was applied with the same parameters as described for the simulated data. For analysis with highly variable genes, 2000 genes were selected using Seurat. The methods were applied in the same way as that with the CD8+ gene signatures.

For the COVID-19 dataset [43], we processed the data as described above. The gene features were selected as the 2000 most variable genes using Seurat. For analysis based on the observed data, the approach was the same as used on the melanoma data. For scINSIGHT analysis, scINSIGHT was applied to the 14 samples treating disease phase as the condition factor. We set *K*_*j*_ = 2 and the best *K* was 13 (selected from {5, 7, 9, 11, 13, 15}). The regularization values were selected as *λ*_1_ = *λ*_2_ = 1. LIGER was applied with nrep=5 and k=13, and the other parameters were set as the default values. Seurat, Harmony, and Scanorama were applied with default parameters. scGen was applied with the same parameters as described for the simulated data. scMerge was applied with kmeansK=rep(3,13), and the other parameters were set as the default values.

For the wound dermis dataset [51], we processed the data using the same approach as applied to the COVID-19 dataset. For scINSIGHT analysis, scINSIGHT was applied to the samples treating treatment/control as the condition factor. We set *K*_*j*_ = 2 and the best *K* was 11 (selected from {5, 7, 9, 11, 13, 15}). The regularization values were selected as *λ*_1_ = *λ*_2_ = 0.1. LIGER was applied with nrep=5 and k=11. The other parameters were set as the default values. Seurat, Harmony, and Scanorama were applied with default parameters. scGen was applied with the same parameters as described for the simulated data. scMerge was applied with kmeansK=c(6,6), and other parameters were set as the default values.

For the mouse retina dataset [30], we processed the data using the same approach as applied to the COVID-19 dataset. For scINSIGHT analysis, scINSIGHT was applied to the two samples treating batches as the condition factor. We set *K*_*j*_ = 2 and the best *K* was 26 (selected from {11, 16, 21, 26, 31}). The regularization values were selected as *λ*_1_ = *λ*_2_ = 1. LIGER was applied with nrep=3 and k=20. The other parameters were set as the default values. Seurat, Harmony, and Scanorama were applied with default parameters. scGen was applied with the same parameters as described for the simulated data, expect for batch size=128. scMerge was applied with kmeansK=c(5,12) and svd k=20, and other parameters were set as the default values.

## Availability of data and materials

The scINSIGHT method has been implemented as an R package, which is available on CRAN (https://cran.r-project.org/web/packages/scINSIGHT/index.html) or at https://github.com/Vivianstats/scINSIGHT. The source code is available at the zenodo repository (DOI: 10.5281/zenodo.5553132) under the GPLv2 license. The melanoma dataset was downloaded from GSE120575 [38]. The COVID-19 dataset was downloaded from FastGenomics (https://beta.fastgenomics.org/p/565003) [43]. The wound dermis dataset was downloaded from GSE112671 [51]. The mouse retina dataset was downloaded from Docker Hub (https://hub.docker.com/repository/docker/jinmiaochenlab/batch-effect-removal-benchmarking) [30].

## Acknowledgments

We thank Dr. Wei-Xing Zong at Rutgers Department of Chemical Biology and the two anonymous reviewers for their insightful suggestions on our study. We also thank Ms. Jie Sheng, a visiting student in the Vivian Li Lab, for the helpful comments on our manuscript. This work was supported by NIH R35GM142702 and R21MH126420, NJ ACTS BERD Mini-Methods Grant (a component of the NIH UL1TR0030117), and Rutgers Busch Biomedical Grant (to W.V.L.).

## Notes

### Competing Interest Statement

The authors have declared no competing interest.

