## Supplementary File for "scINSIGHT for interpreting single-cell gene expression from biologically heterogeneous data"

### Supplementary Figures and Table

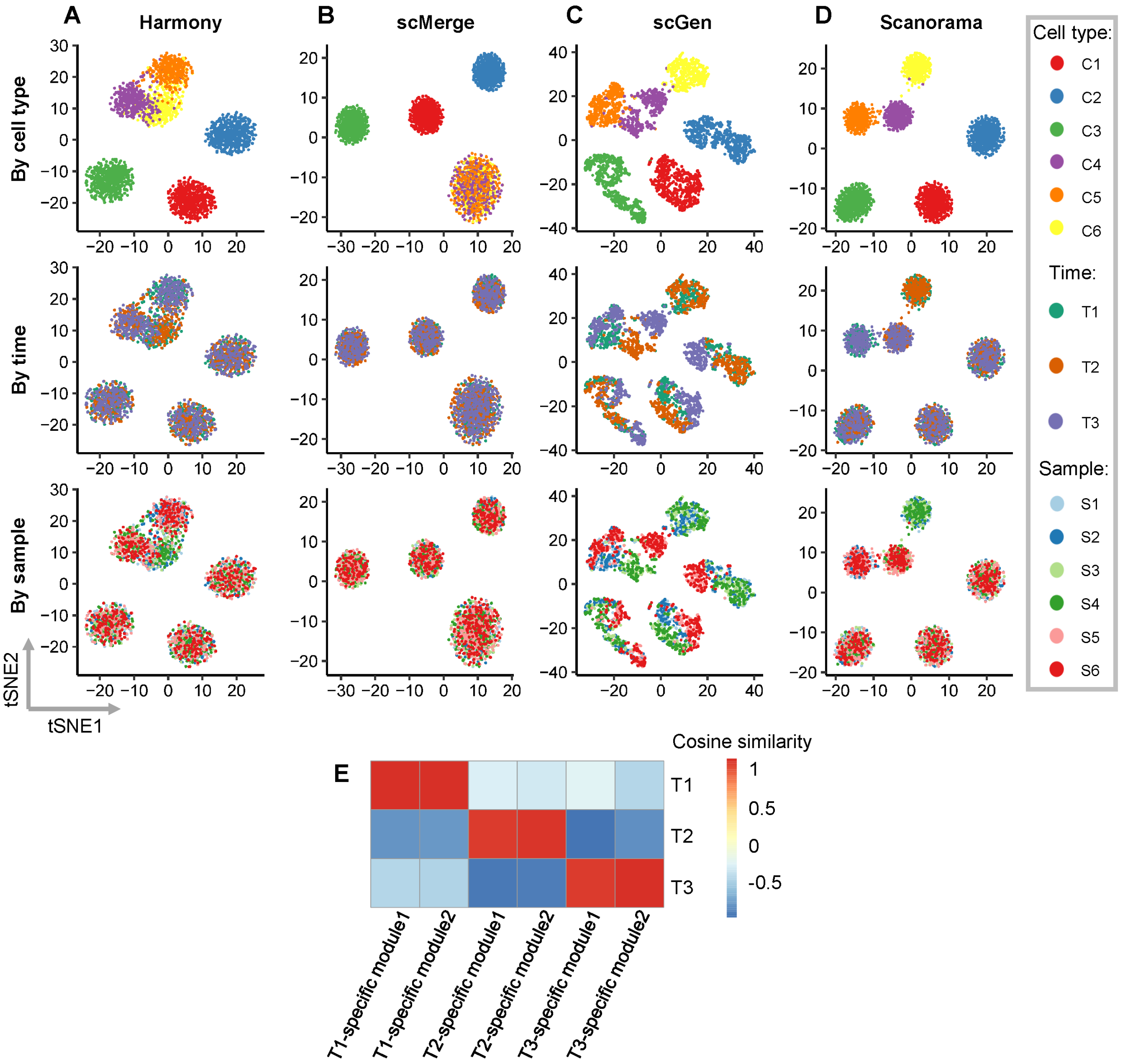


**Figure S1:** Comparison of integrated data in the simulation study. **A-D**: tSNE plots of simulated cells based on the integrated data by Harmony (**A**), scMerge (**B**), scGen (**C**) and Scanorama (**D**). For each method, three tSNE plots colored by cell type, time point, or sample index are displayed. **E**: Cosine similarity between true time-point effects and membership vectors of condition-specific gene modules obtained by scINSIGHT.

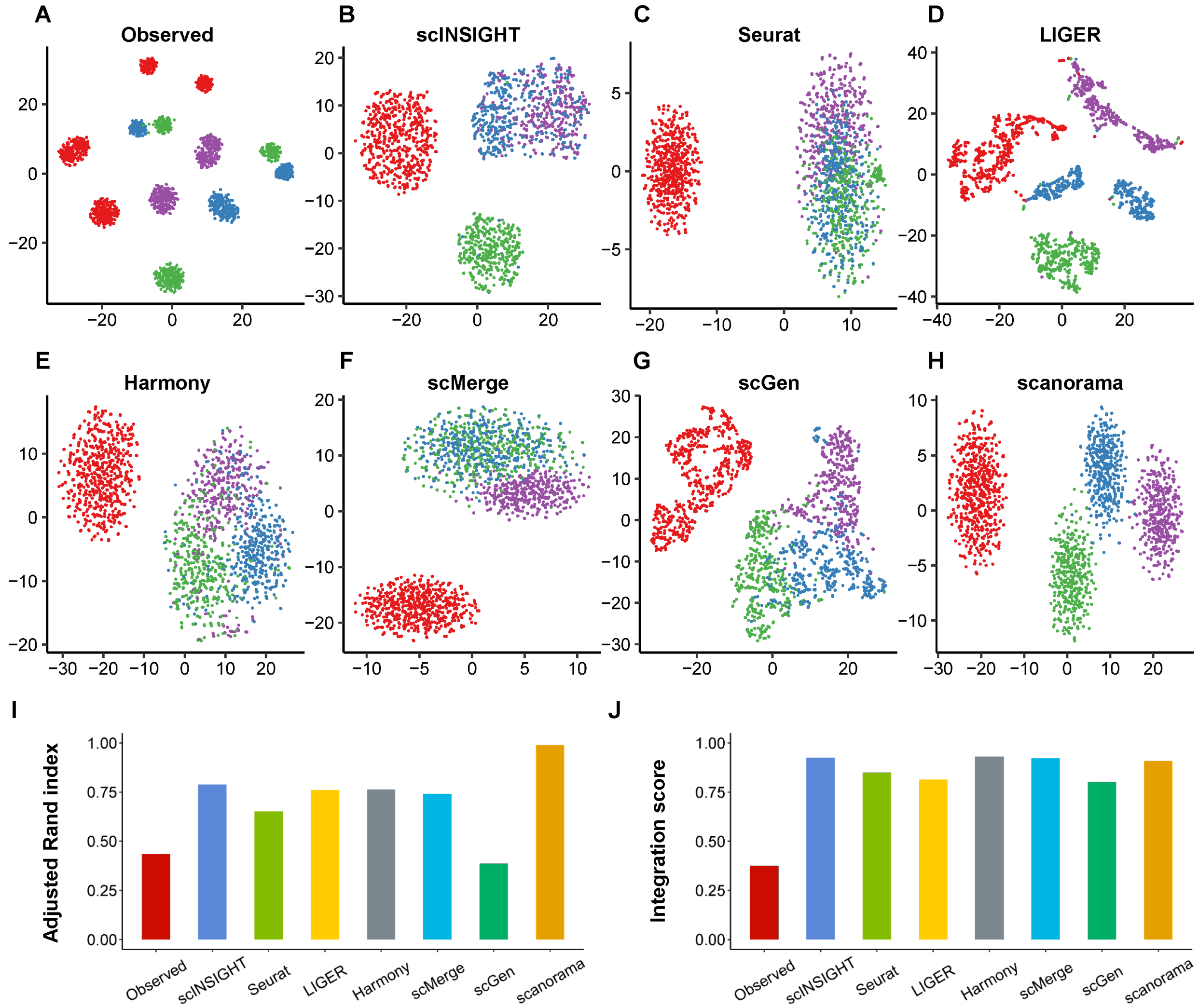


**Figure S2:** Comparison of observed and integrated data in the simulation study (variant 1). **A-H**: tSNE plots of simulated cells based on the observed (unintegrated) data (**A**) and integrated data by scINSIGHT (**B**), Seurat (**C**), LIGER (**D**), Harmony (**E**), scMerge (**F**), scGen (**G**) and Scanorama (**H**). For each method, tSNE plot colored by cell type is displayed. **I**: Adjusted Rand index calculated using clusters identified from the observed or integrated data. **J**: Integration scores of the observed and integrated data.

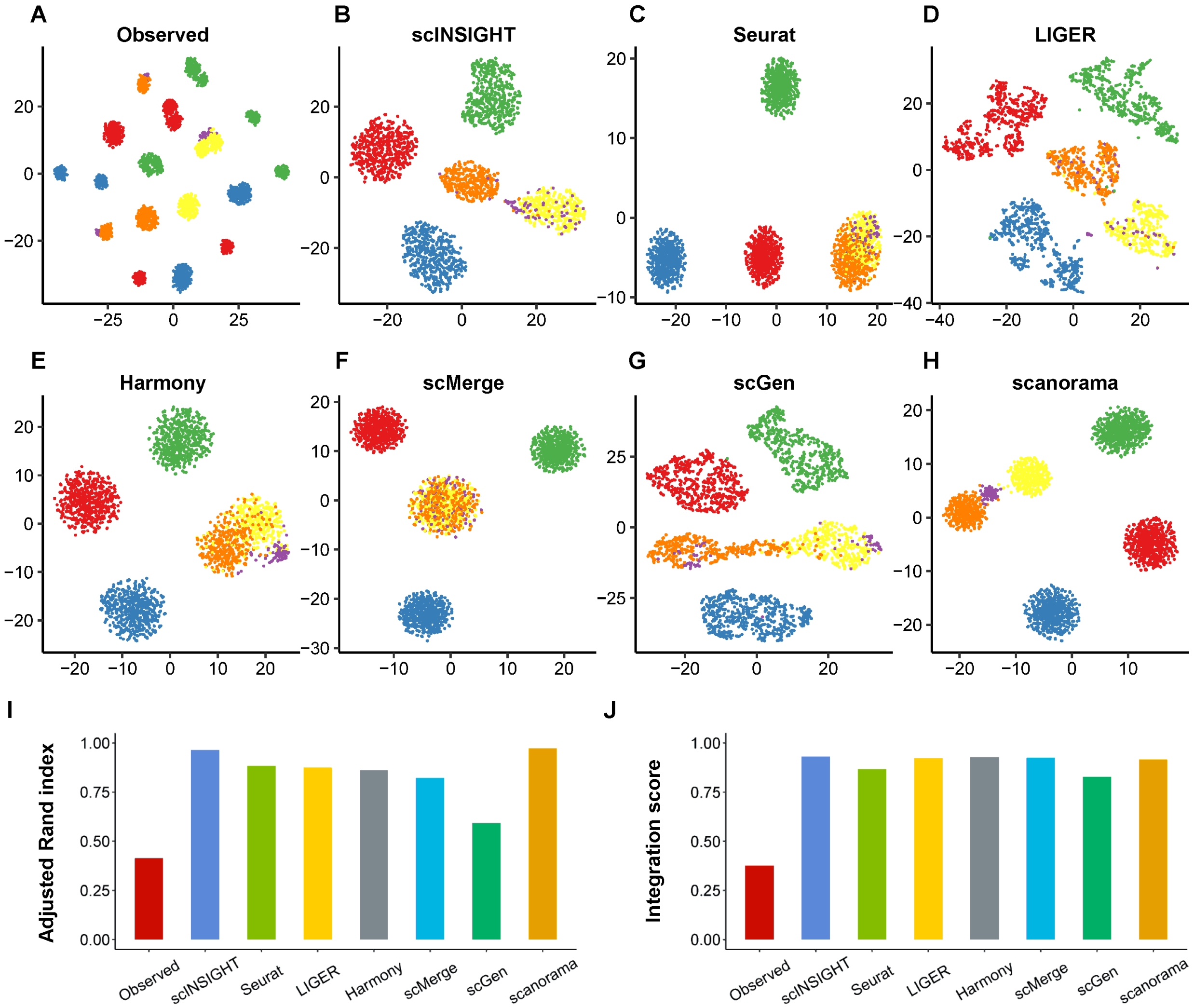


**Figure S3:** Comparison of observed and integrated data in the simulation study (variant 3). **A-H**: tSNE plots of simulated cells based on the observed (unintegrated) data (**A**) and integrated data by scINSIGHT (**B**), Seurat (**C**), LIGER (**D**), Harmony (**E**), scMerge (**F**), scGen (**G**) and Scanorama (**H**). For each method, tSNE plot colored by cell type is displayed. **I**: Adjusted Rand index calculated using clusters identified from the observed or integrated data. **J**: Integration scores of the observed and integrated data.

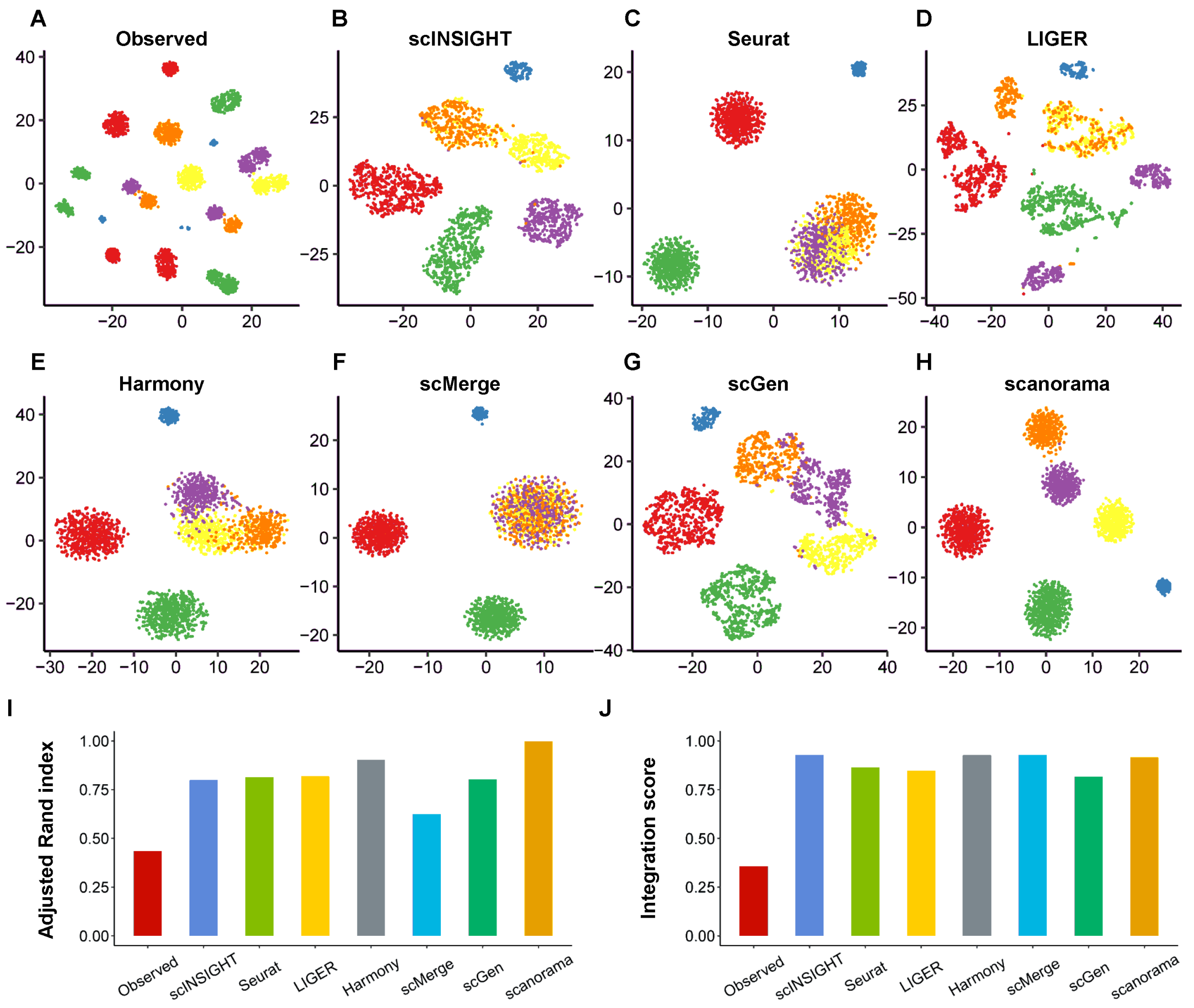


**Figure S4:** Comparison of observed and integrated data in the simulation study (variant 2). **A-H**: tSNE plots of simulated cells based on the observed (unintegrated) data (**A**) and integrated data by scINSIGHT (**B**), Seurat (**C**), LIGER (**D**), Harmony (**E**), scMerge (**F**), scGen (**G**) and Scanorama (**H**). For each method, tSNE plot colored by cell type is displayed. **I**: Adjusted Rand index calculated using clusters identified from the observed or integrated data. **J**: Integration scores of the observed and integrated data.

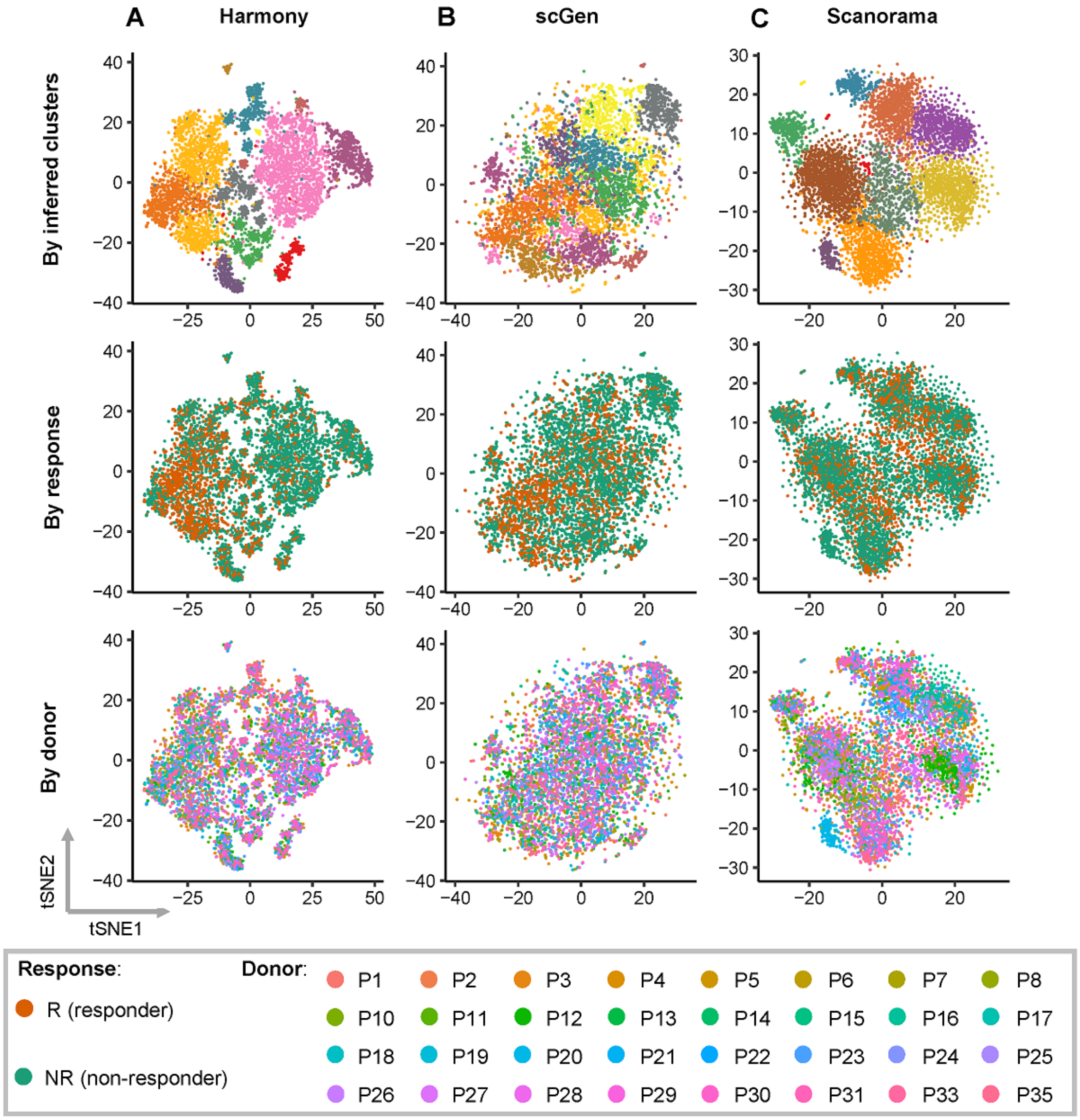


**Figure S5:** tSNE plots of CD8+ T cells from melanoma patients based on the integrated data by Harmony (**A**), scGen (**B**) and Scanorama (**C**). For each method, three tSNE plots colored by inferred cell cluster, NR/R condition, or donor index are displayed.

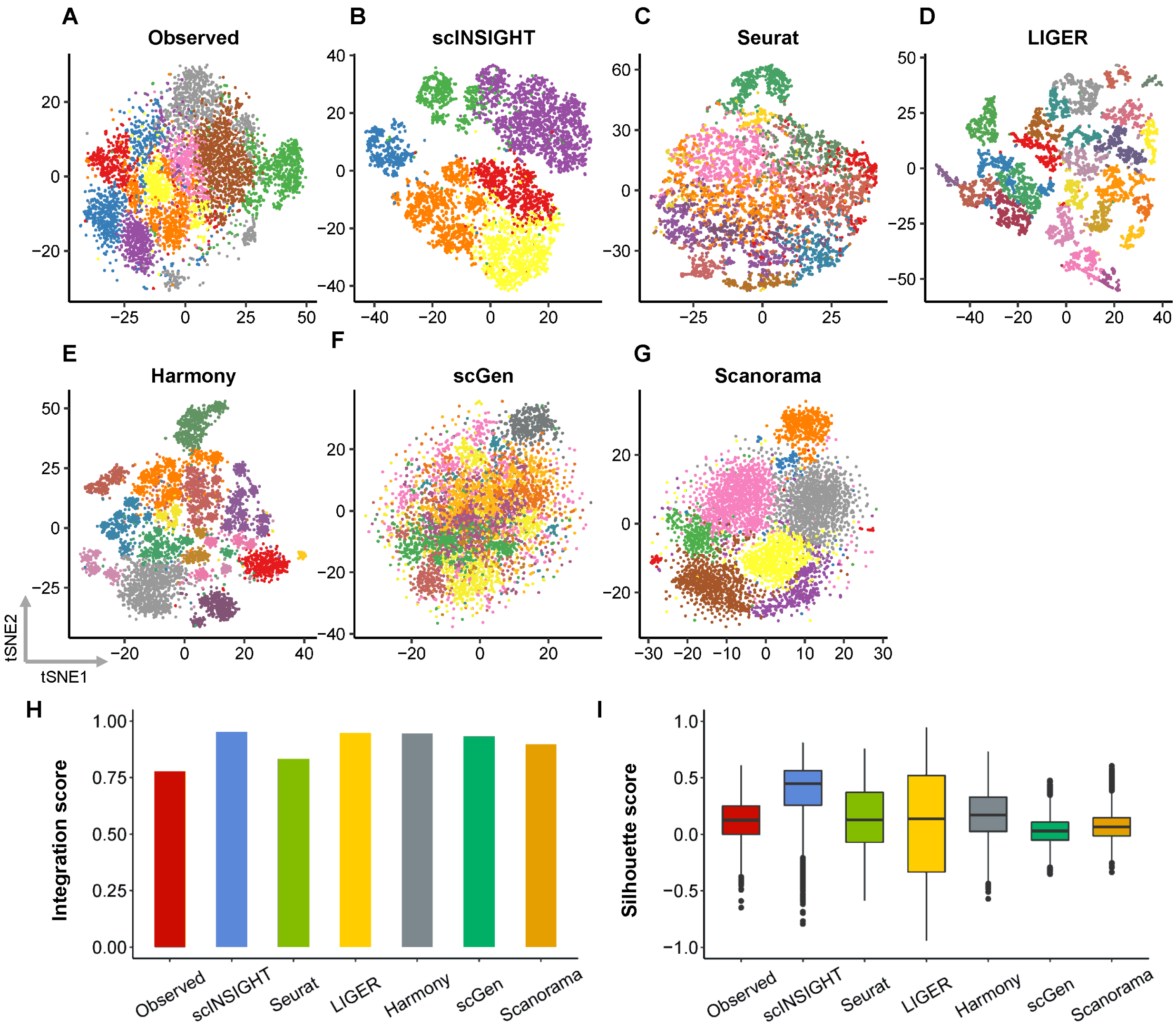


**Figure S6:** Comparison of observed and integrated data in the melanoma study (using highly variable genes). **A-G**: tSNE plots of simulated cells based on the observed (unintegrated) data (**A**) and integrated data by scINSIGHT (**B**), Seurat (**C**), LIGER (**D**), Harmony (**E**), scGen (**F**) and Scanorama (**G**). Cells are colored by inferred cell cluster. **H**: Adjusted Rand index calculated using clusters identified from the observed or integrated data. **I**: Integration scores of the observed and integrated data.

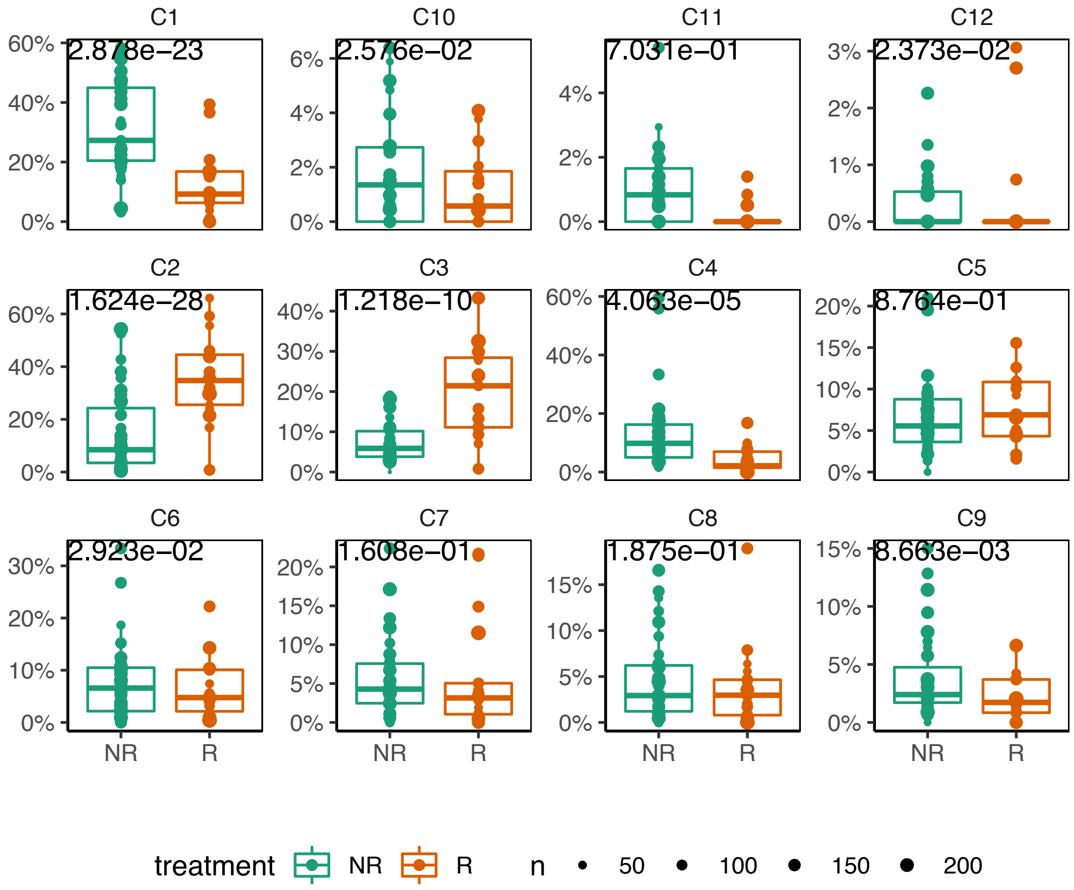


**Figure S7:** Percentage of the clusters identified by Harmony. $P$-values indicate significance of association between cluster proportion and response, and were calculated by ANOVA for the logistic regression model.

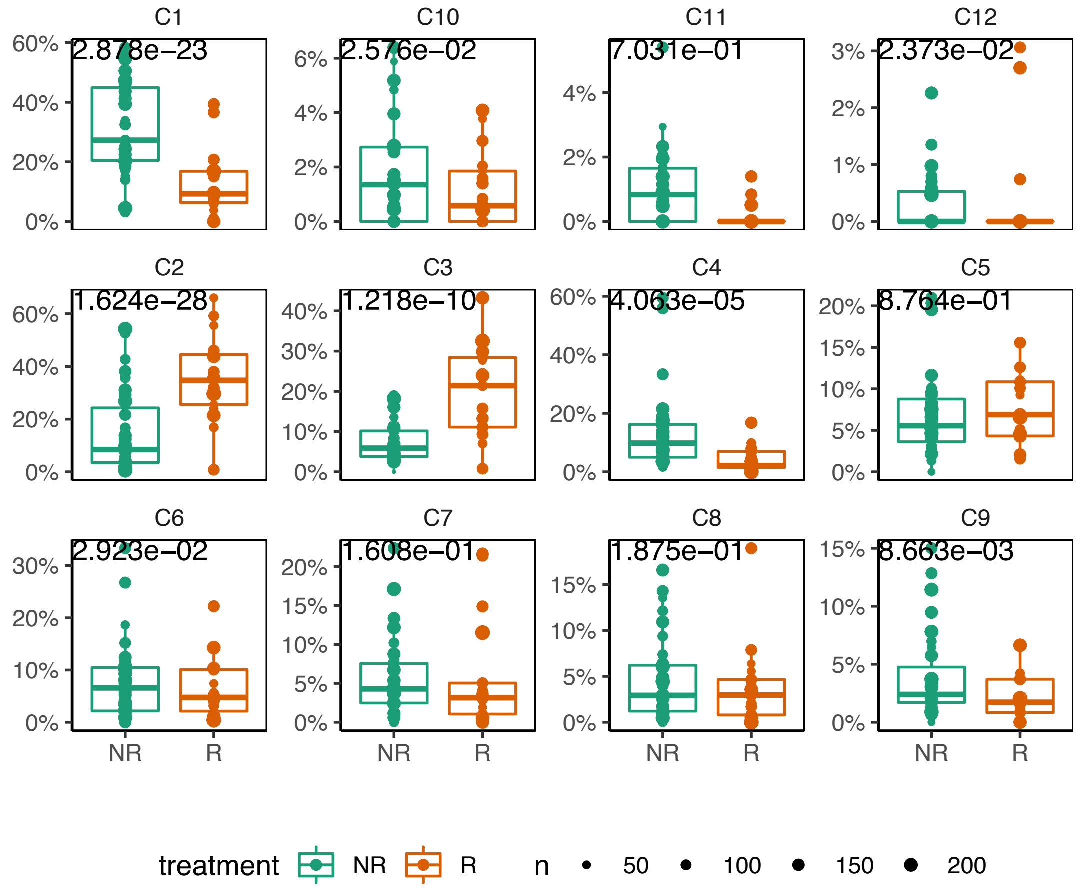


**Figure S8:** Percentage of the clusters identified by scGen. $P$-values indicate significance of association between cluster proportion and response, and were calculated by ANOVA for the logistic regression model.

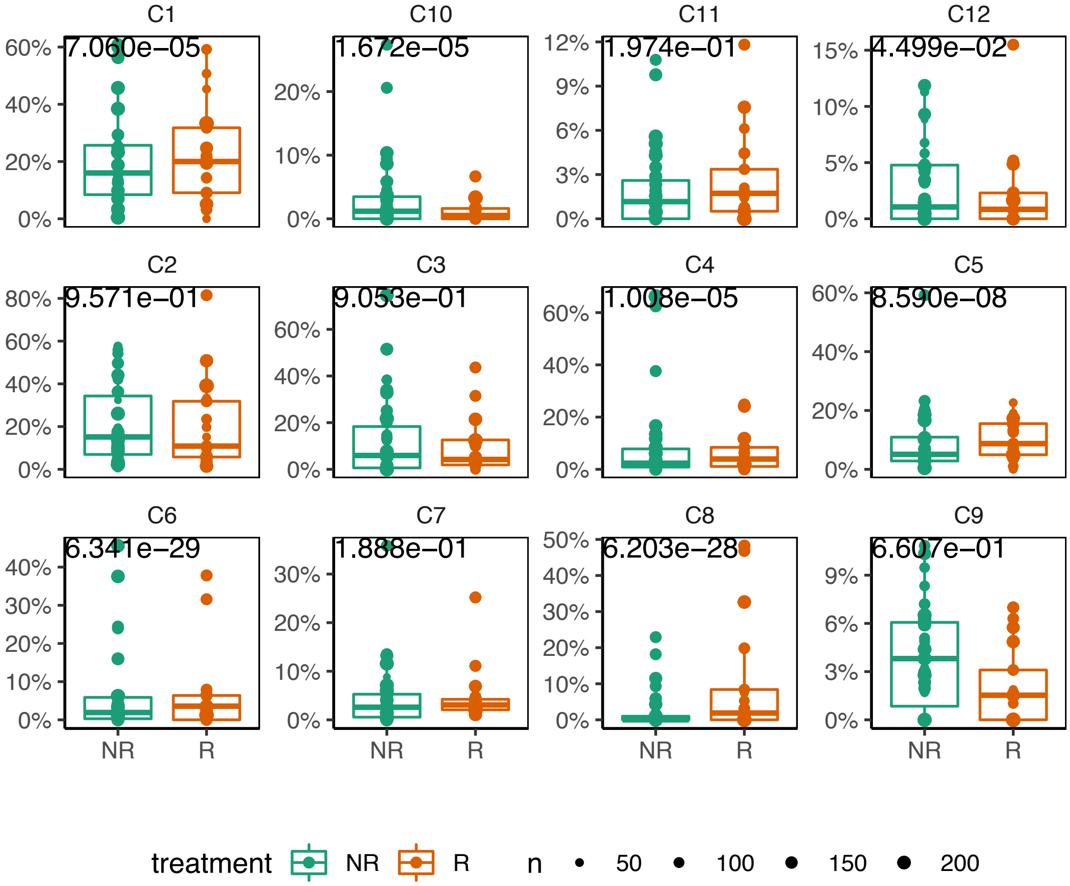


**Figure S9:** Percentage of the clusters identified by Seurat. $P$-values indicate significance of association between cluster proportion and response, and were calculated by ANOVA for the logistic regression model.

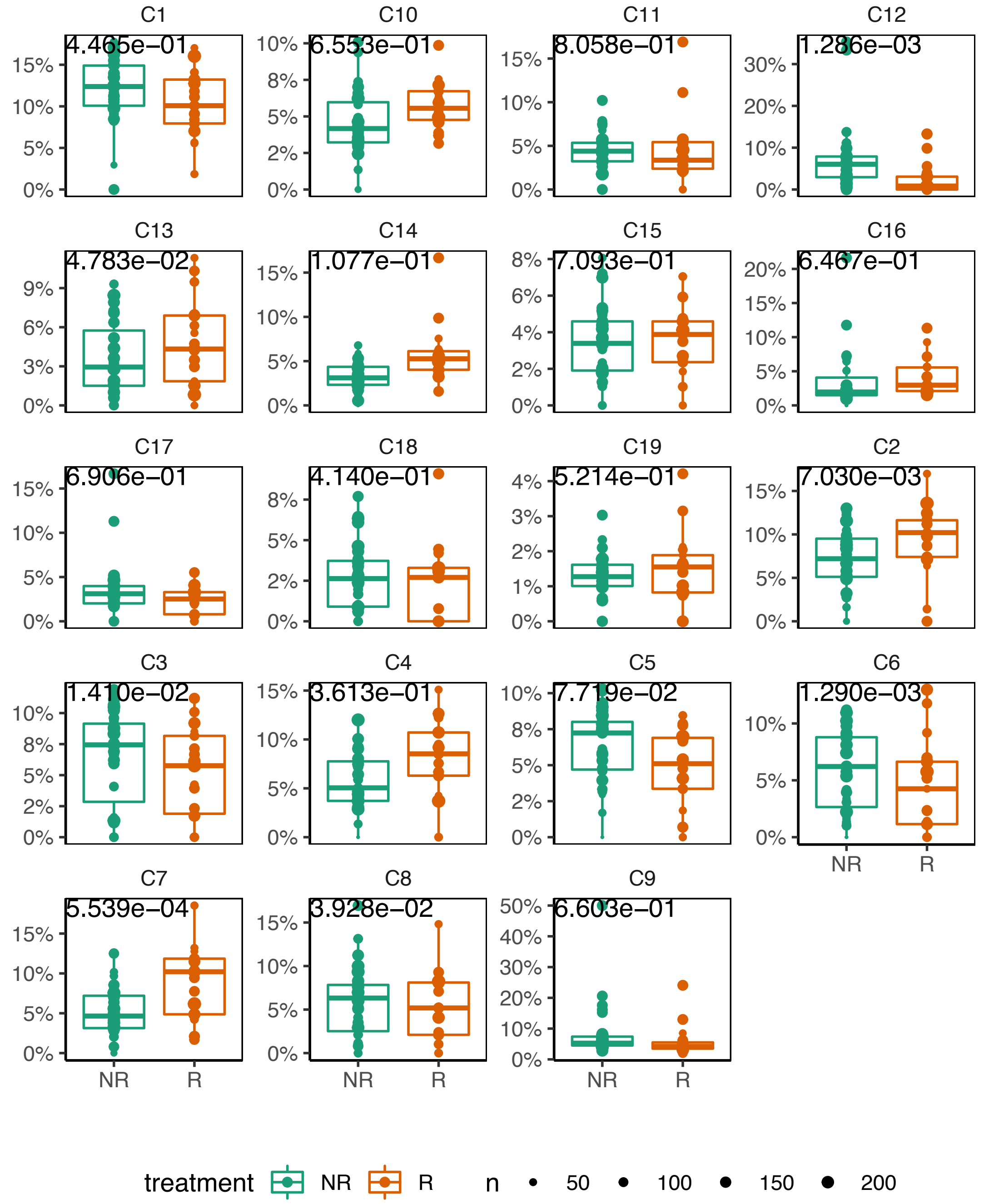


**Figure S10:** Percentage of the clusters identified by LIGER. $P$-values indicate significance of association between cluster proportion and response, and were calculated by ANOVA for the logistic regression model.

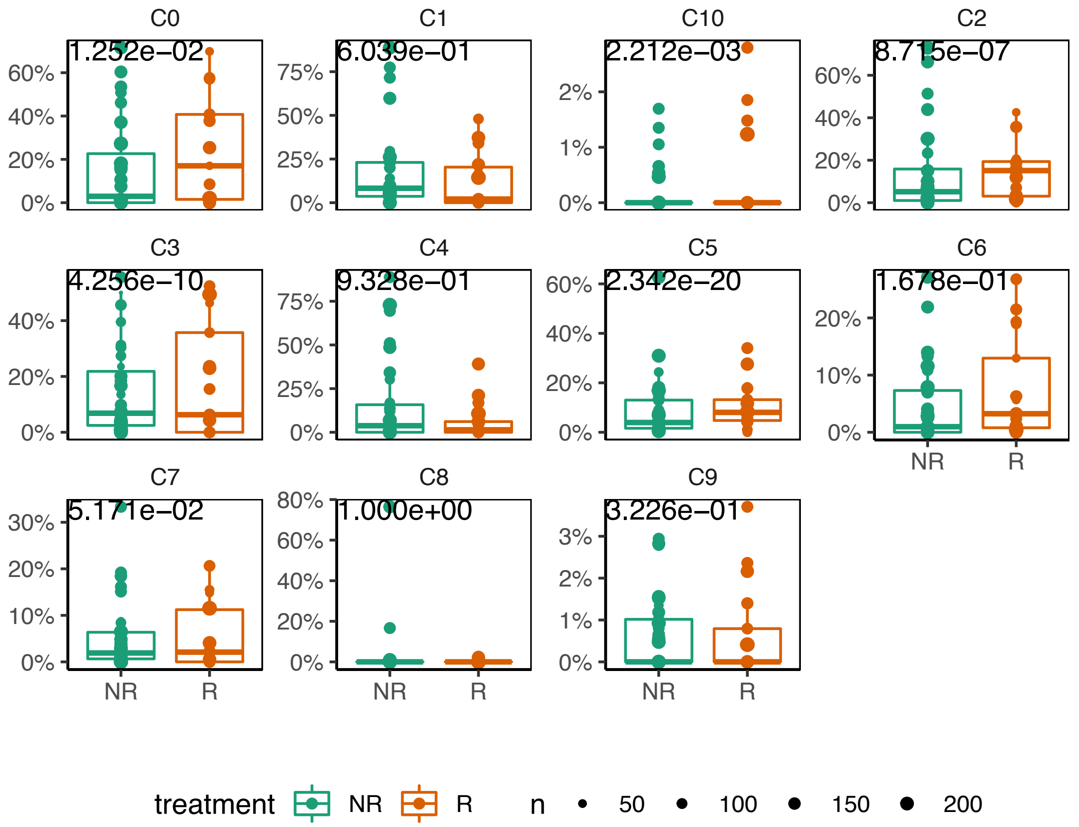


**Figure S11:** Percentage of the clusters identified by Scanorama. $P$-values indicate significance of association between cluster proportion and response, and were calculated by ANOVA for the logistic regression model.

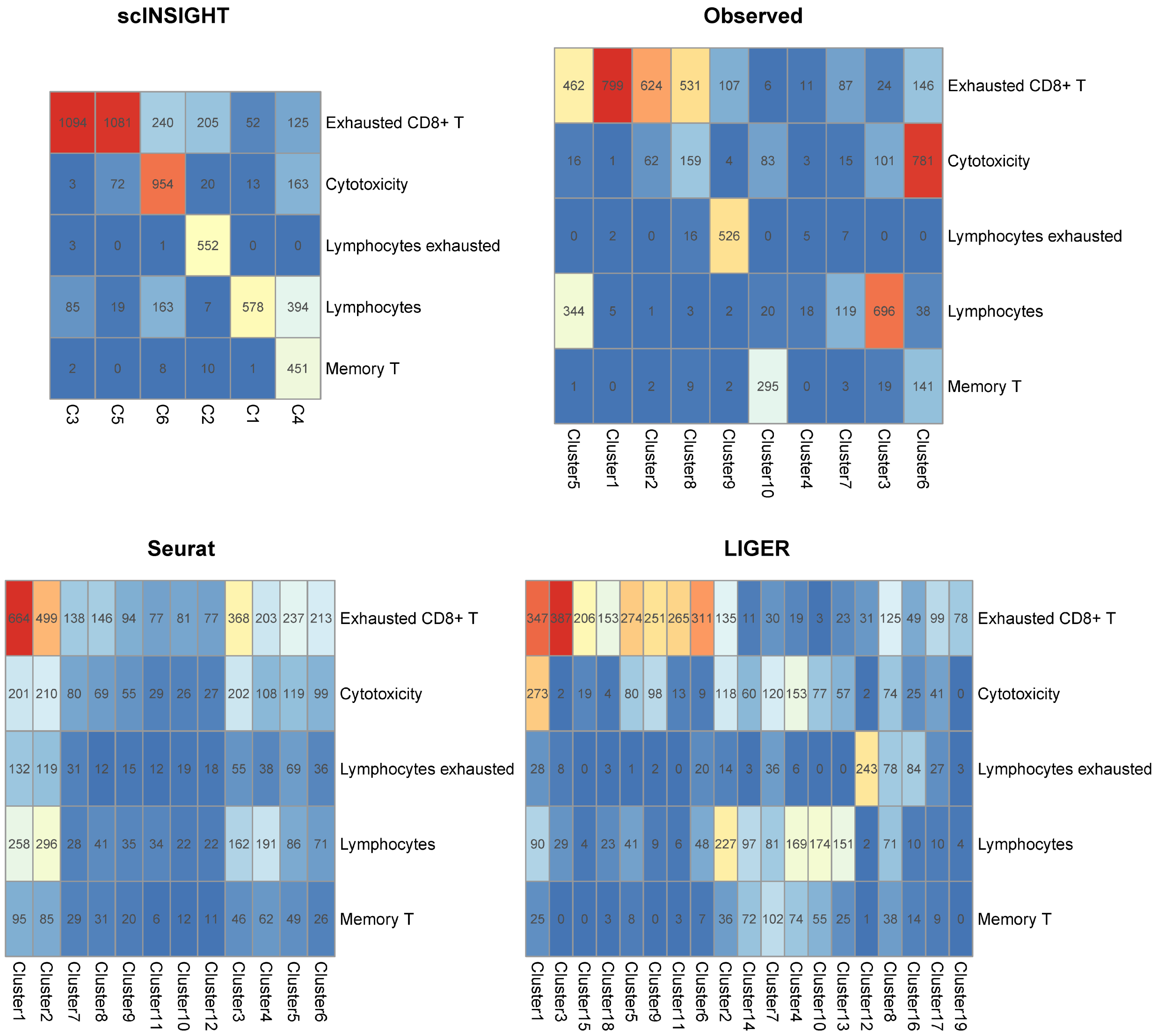


**Figure S12:** Contingency tables of cell type annotations presented in Sade-Feldman et al (original publication) and computationally inferred cell clusters based on observed data and integrated data by scINSIGHT, Seurat, and LIGER.

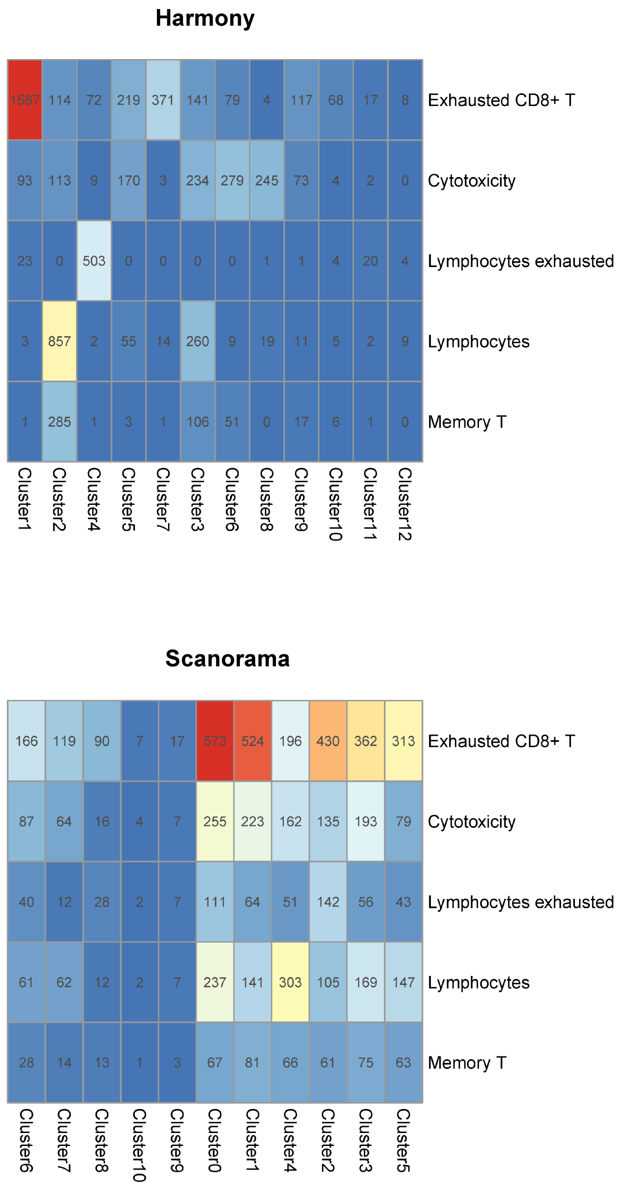


**Figure S13:** Contingency tables of cell type annotations presented in Sade-Feldman et al (original publication) and computationally inferred cell clusters based on observed data and integrated data by Harmony, scGen, and Scanorama.

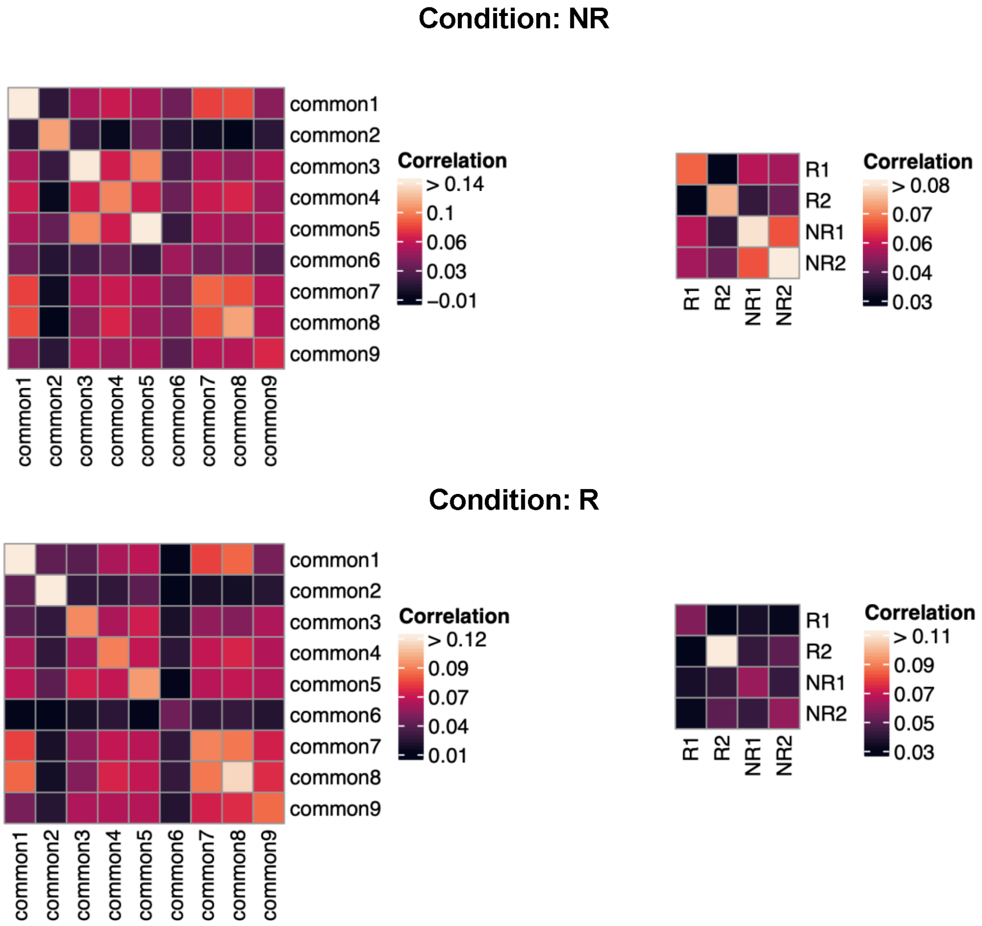


**Figure S14:** Within-module and between-module co-expression in the melanoma dataset. For each gene module, we selected the top 25 genes with the largest coefficients, and used these genes to calculate within-module and between-module co-expression scores. The within-module co-expression in one condition was calculated as the average pair-wise Spearman's correlation of the top genes in that module, using all cells from that condition. The between-module co-expression in one condition was calculated as the average Spearman's correlation between top genes in the two modules, using all cells from that condition.

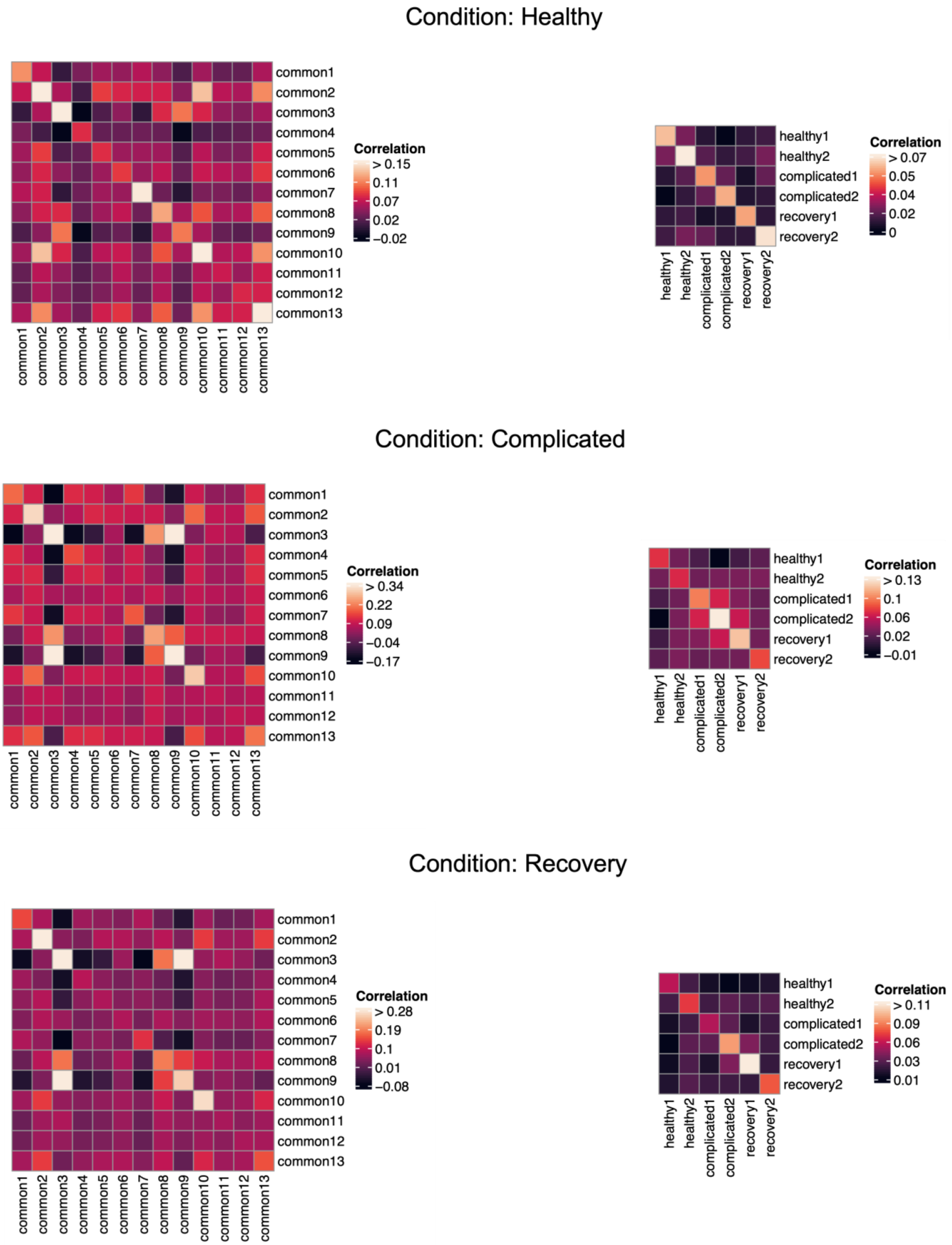


**Figure S15:** Within-module and between-module co-expression in the COVID-19 dataset.

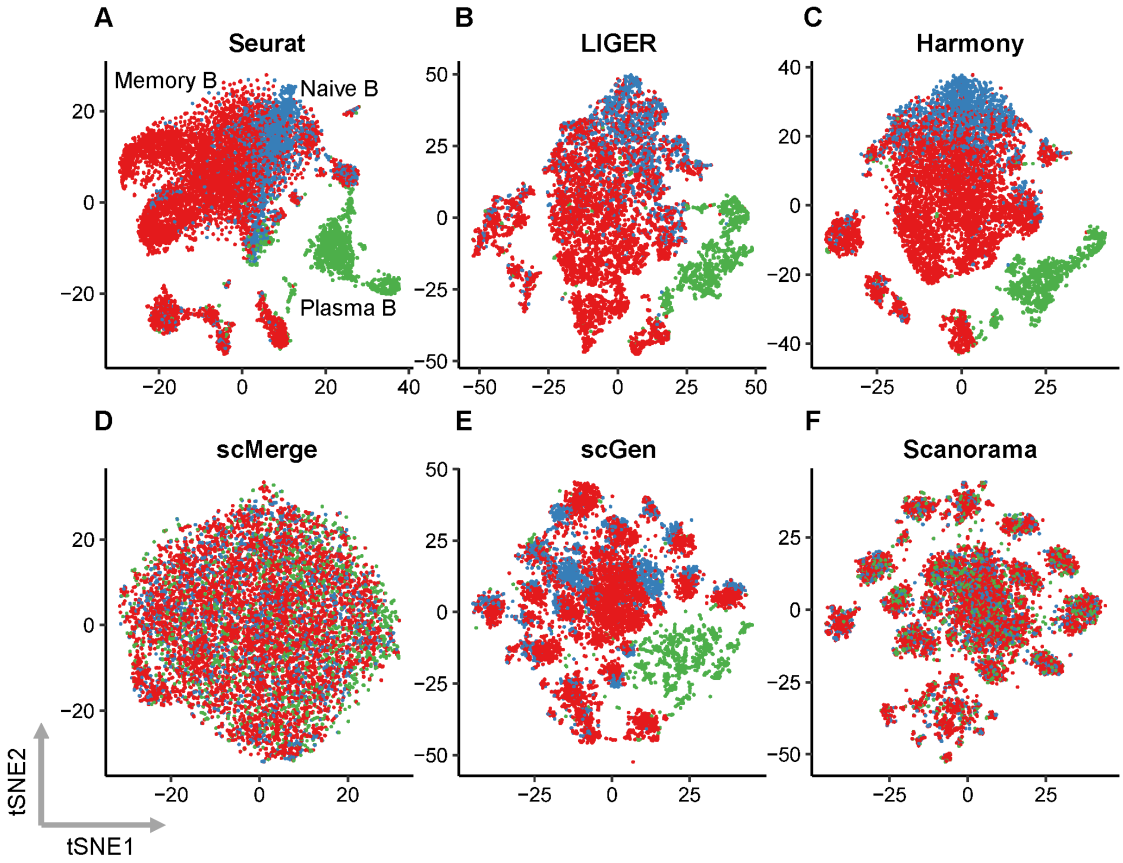


**Figure S16:** tSNE plots of B cells based on integrated data by Seurat (**A**), LIGER (**B**), Harmony (**C**), scMerge (**D**), scGen (**E**) and Scanorama (**F**). Cells are colored by classified cell type (by SingleR).

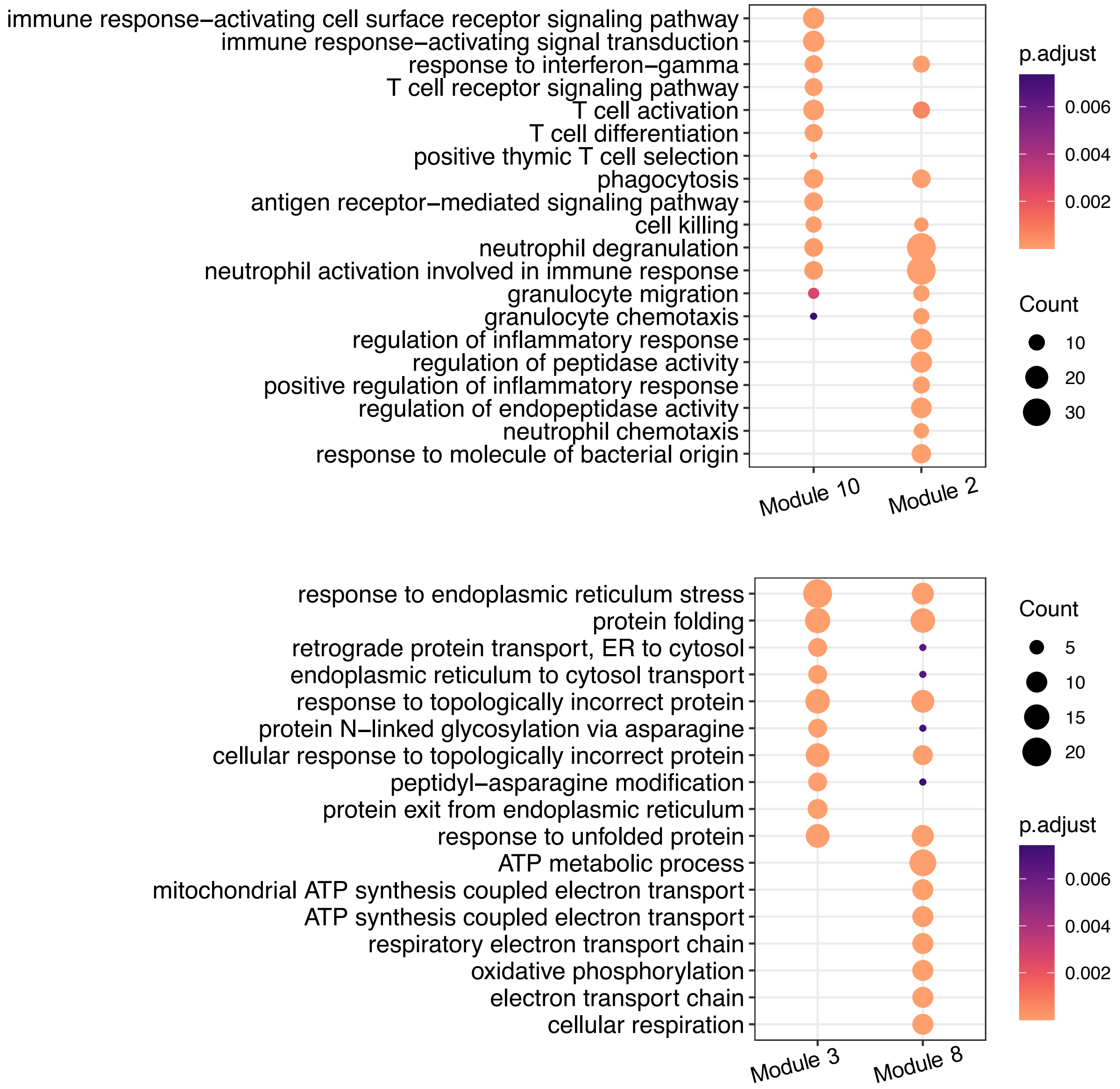


**Figure S17:** Top enriched GO terms in common gene modules (10, 2, 3, and 8) identified by scINSIGHT.

### Condition: Hh activation

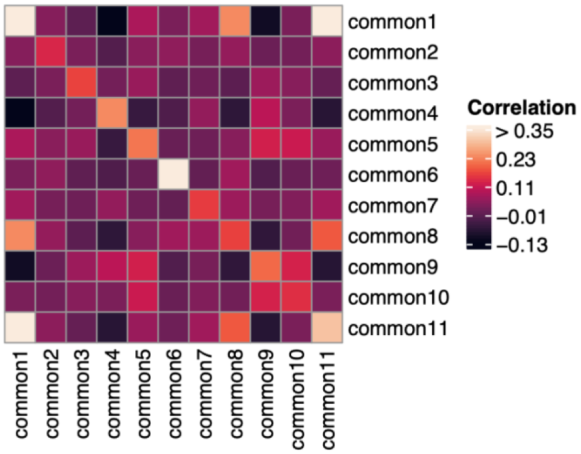


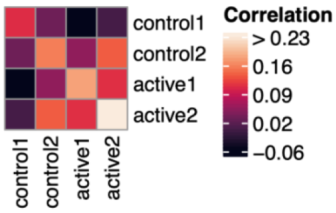


### Condition: Control

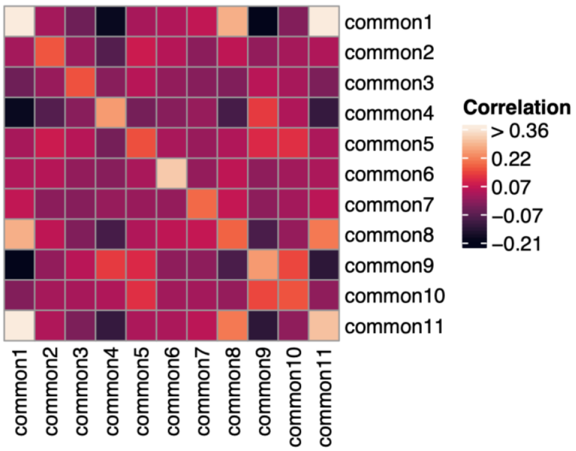


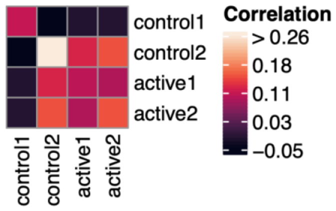


**Figure S18:** Within-module and between-module co-expression in the Wound healing dataset.

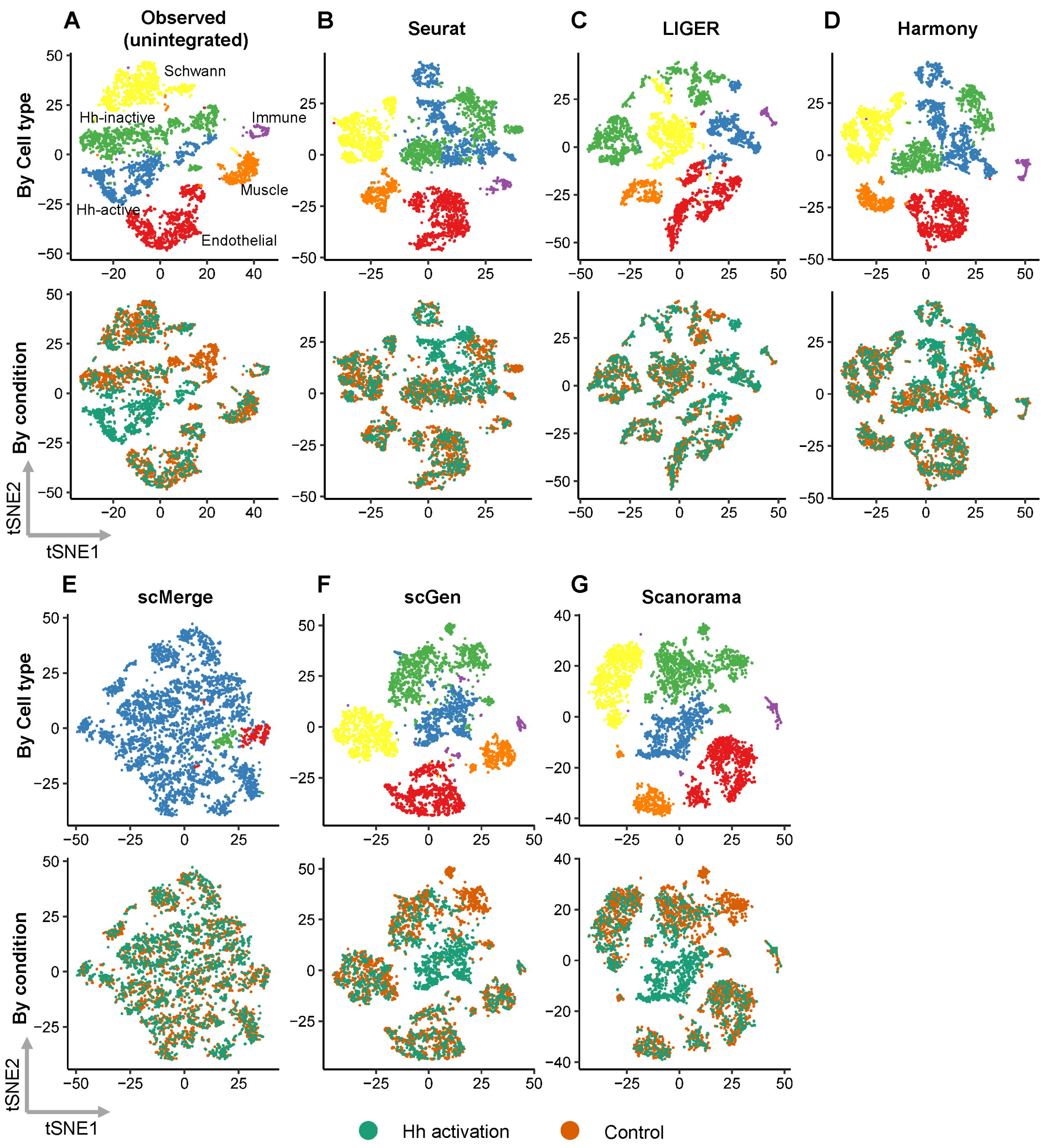


**Figure S19:** tSNE plots of dermal cells based on observed data (**A**) and integrated data by Seurat (**B**), LIGER (**C**), Harmony (**D**), scMerge (**E**), scGen (**F**), and Scanorama (**G**).

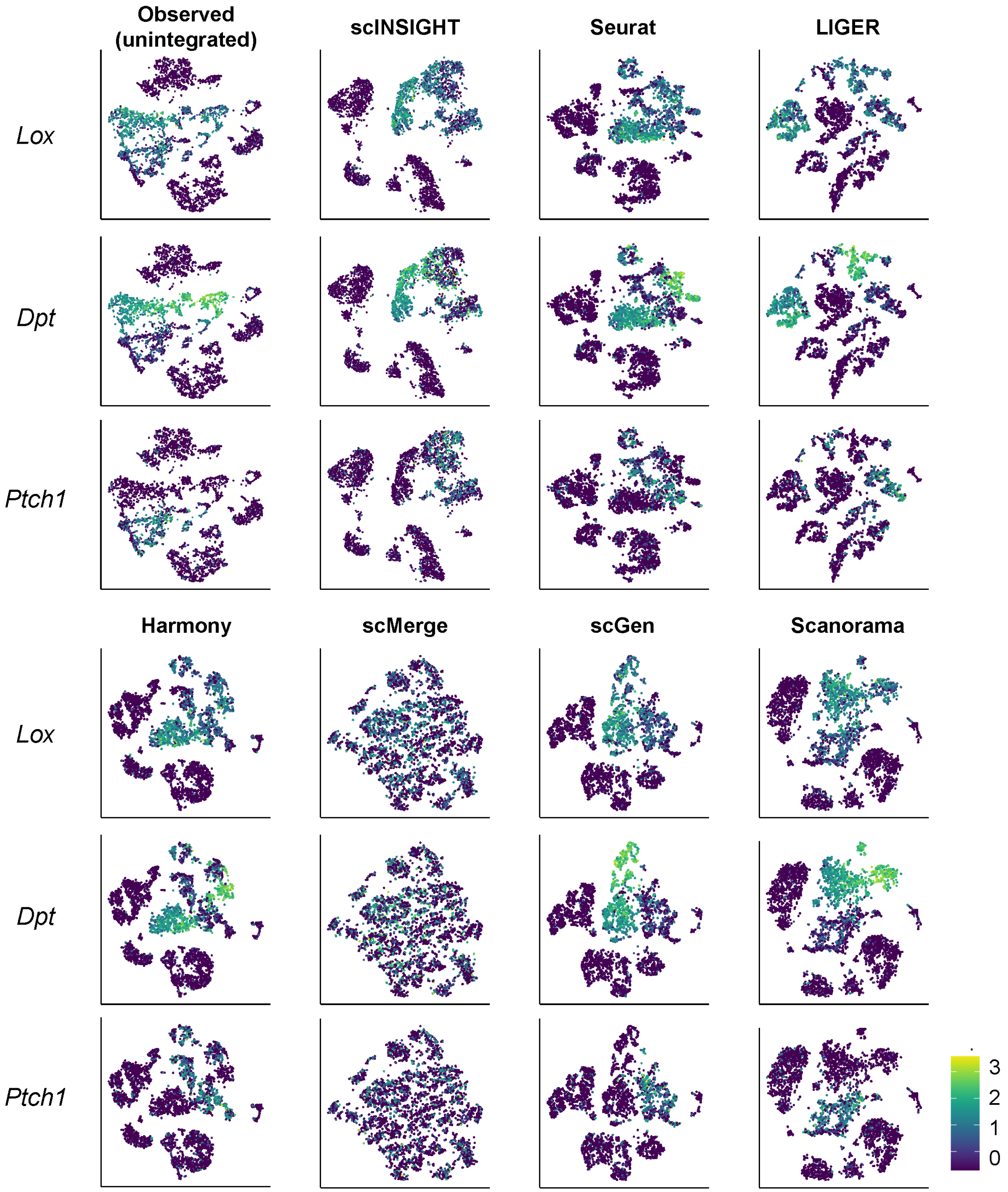


**Figure S20:** tSNE plots of observed and integrated data colored by scaled expression of fibroblast signatures (*Lox*, *Dpt*, and *Ptch1*).

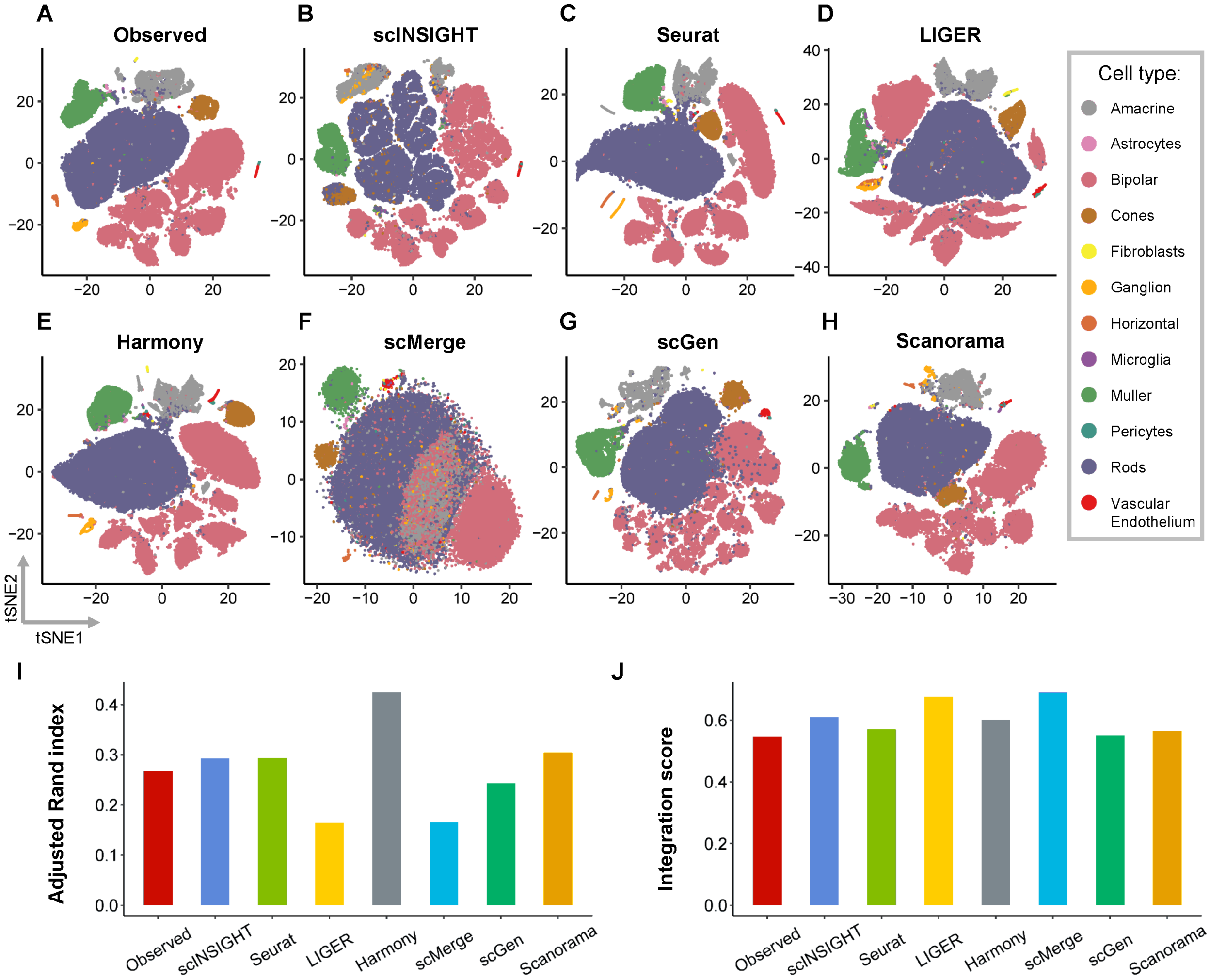


**Figure S21:** Comparison of observed and integrated data in the mouse retina study. **A-H**: tSNE plots of single cells based on the observed (unintegrated) data (**A**) and integrated data by scINSIGHT (**B**), Seurat (**C**), LIGER (**D**), Harmony (**E**), scMerge (**F**), scGen (**G**), and Scanorama (**H**). Cells are colored by cell type. **I**: Adjusted Rand index calculated using clusters identified from the observed or integrated data. **J**: Integration scores of the observed and integrated data.

**Table S1:** Running time and memory usage of scINSIGHT and the other six integration methods. For scINSIGHT, the recorded time on the top was time used to run scINSIGHT with different values of $K$, and the recorded time on the bottom (in the parentheses) was time used to select regularization parameters.

| Dataset | | scINSIGHT | Seurat | LIGER | Harmony | scMerge | scGen | Scanorama |
|---|---|---|---|---|---|---|---|---|
| Simulation<br>3000 cells<br>6 samples<br>3 conditions | Memory | 5.91G | 7.96G | 1.49G | 1.10G | 2.37G | 12.06G | 6.72G |
| | Time | 12479.10s<br>(3751.98s) | 148.37s | 524.57s | 3.35s | 528.60s | 87.59s | 4.43s |
| Melanoma<br>6350 cells<br>48 samples<br>2 conditions | Memory | 17.70G | 10.45G | 9.75G | 11.85G | | 23.06G | 14.45G |
| | Time | 6945.26s<br>(3308.20s) | 105.68s | 340.34s | 41.67s | | 296.70s | 8.01s |
| COVID-19<br>9741 cells<br>13 samples<br>3 conditions | Memory | 14.52G | 13.25G | 4.00G | 5.16G | 5.88G | 18.76G | 11.65G |
| | Time | 6753.42s<br>(3733.60s) | 702.68s | 697.63s | 32.47s | 209.16s | 494.31s | 9.55s |
| Wound healing<br>4680 cells<br>2 samples<br>2 conditions | Memory | 7.63G | 3.15G | 2.26G | 3.50G | 3.61G | 16.16G | 9.75G |
| | Time | 2385.18s<br>(849.12s) | 60.65s | 458.36s | 16.75s | 78.32s | 207.95s | 1.92s |
| Mouse retina<br>71638 cells<br>2 samples<br>2 conditions | Memory | 49.5G | 23.75G | 15.55G | 16.25G | 54.65G | 46.36G | 29.55G |
| | Time | 80221.78s<br>(50878.43s) | 6381.78s | 14910.96s | 140.58s | 1532.17s | 2457.38s | 55.82s |
